# Effect of Stapling on the Thermodynamics of mdm2-p53 Binding

**DOI:** 10.1101/2020.12.28.424518

**Authors:** Atanu Maity, Asha Rani Choudhury, Rajarshi Chakrabarti

## Abstract

Protein-protein interaction (PPI) is one of the key regulatory features to drive biomolecular processes and hence is targeted for designing therapeutics against diseases. Small peptides are a new and emerging class of therapeutics owing to their high specificity and low toxicity. For achieving efficient targeting of the PPI, amino acid side chains are often stapled together resulting in the rigidification of these peptides. Exploring the scope of these peptides demands a comprehensive understanding of their working principle. In this work, two stapled p53 peptides have been considered to delineate their binding mechanism with mdm2 using computational approaches. Addition of stapling protects the secondary structure of the peptides even in the case of thermal and chemical denaturation. Although the introduction of a stapling agent increases the hydrophobicity of the peptide, the enthalpic stabilization decreases. This is overcome by the lowering of the entropic penalty and the overall binding affinity improves. The mechanistic insights into the benefit of peptide stapling can be adopted for further improvement of peptide therapeutics.

## Introduction

Proteins are part of several important biomolecular processes like enzyme catalysis, signal transduction, etc. and they regulate these processes at different stages. Regulation of these processes by protein is governed by specific protein-protein interactions.^1^ In the diseased condition, these sophisticated protein-protein interactions are compromised, inhibited, or dis-regulated.^2^ The strategy to design ways to fight diseases involves restoring those interactions by using some chemical moieties which can mimic one of the original partners in the protein-protein interaction.^3^ Different types of molecules are used as therapeutics^4^ e.g. small organic molecules,^5^ peptides,^6^ biologics such as growth factor, antibodies^7^ etc. The small molecule inhibitors (having molecular weight below 500 Da) and the biologics (having molecular weight above 5000 Da) form the two ends of the molecular inhibitors in terms of molecular weight.^8^ Due to their size, the small molecule inhibitors have high oral bioavailability and permeability. However, sometimes they suffer from limited target selectivity due to their chemical nature. On the other hand, the biologics are very target specific due to their larger interaction area with their target and shows high affinity. Unfortunately because of their size, they suffer from low membrane permeability and metabolic stability. Considering the advantages and disadvantages of these two classes, a third class of therapeutic having a molecular weight in between small molecules and biologics is emerging as a potential improvement.^9,10^ These are orally bioavailable, cell-permeable and have lower manufacturing costs.^11^ Members of this class are mainly peptides derived from naturally occurring proteins and their modified or engineered versions. The design of these bio-active molecules has immensely improved in the last decade to increase their efficiency by introducing chemical modifications to their conformation. One such chemical modification is known as ‘cross-linking’ or ‘stapling’ and the modified peptides are known as ‘stapled peptides’.^12,13^ The side chains of some suitably positioned amino acids are replaced by a variety of chemical moieties, and joined together using ring-closing metathesis. This puts a restrain to the backbone conformations of the peptide which results in a loss of its conformational flexibility and the entropic penalty during complexation is reduced. The design of a stapled peptide can be modulated by controlling two factors - (i) nature of stapling agent and (ii) stapling position. The most used stapling agent is a simple aliphatic hydrocarbon chain although several other modifications are also used like lactam,^14^ aromatic hydrocarbon^15^ etc. The most optimized stapling positions are *i*−*i*+4 and *i*−*i*+7 where the amino acid at the *i*th position is cross-linked with the amino acid at (*i*+4)th and (*i*+7)th position respectively. There are some example of *i*−*i*+3^16^ and *i*−*i*+11^17^ stapled peptides also. Examples of proteins targeted by stapled peptide are mdm2/mdmx,^18^ anti-apoptotic Bcl-2 family proteins,^19^ estrogen receptor^20^ etc. Amongst these, the ones targeting mdm2 are very important in the context of designing therapeutics.

The p53 tumor suppressor is a crucial modulator of growth arrest and apoptosis in response to cellular damage and stress.^21^ For example, in response to DNA damage, it controls the growth arrest to allow DNA repair or propagates apoptosis. These prevent the proliferation of damaged cells and transfer of lethal mutations to the next generation of cells.^22^ In normal cellular condition, p53 is present in the cell in very low concentration and their tumor suppressor function is antagonized by mdm2.^23^ Interestingly mdm2 is overexpressed in cells for different types of tumors.^24,25^ The cancerous cells produce more mdm2 to inactivate p53 to ensure their growth. Designing compounds that can prevent the interaction between p53 and mdm2 is therefore an attractive strategy to restore p53 tumor-suppressor activity in tumors. There are approaches to design both small-molecule inhibitor^26,27^ as well as peptide-based inhibitor,^28^ ‘stapled peptide’ in particular. Significant numbers of stapled peptides have been proposed and their mdm2-bound structures have been solved using X-ray crystallography^29^ and NMR spectroscopy,^16^ yet the success in the clinical trials is not very promising. For example, a stapled peptide developed by Chang *et al*. as a dual inhibitor of mdm2 and mdmx has passed the phase-Ib clinical trial.^30^ This limitation of the designed peptides demands a detailed understanding of the binding mechanism of stapled peptides with mdm2. The structural details available from crystallographic structures are not sufficient as it does not capture the dynamic aspect of the protein-peptide interaction however, the structures obtained using NMR spectroscopy often provide useful information about the dynamics.^16^ In this context, a computational approach combining molecular modeling and molecular dynamics simulation can provide significant insight. Significant progress has been made by several research groups in this direction which includes comparisons of stapled and unstapled peptide binding, benchmarks of stapling position on the peptide from the calculated thermodynamic properties,^31–34^ comparison of performances of force fields to model stapled peptide,^35^ and development of new simulation methodology which can address the kinetics of binding.^36^ Most of these studies address the changes in p53 conformation inside the binding pocket upon addition of the stapling agent. Though that gives a considerable picture of the binding process, the whole picture will not be clear without considering the free state conformations. While going from its free state (having large numbers of possible conformations) to the mdm2-bound state (having effectively reduced numbers of conformations), the p53 peptide will undergo a large conformational change. This transformation from high entropy state to low entropy state will cost entropic penalty to the overall binding energy. Since stapling agents introduce positional restraint in the peptide that can be expected to influence the free sate and hence the entropic penalty value. Therefore it is equally important to compare the free-state conformation of p53 and stapled p53.

The binding of mdm2 with three peptides p53, p53 with an aliphatic hydrocarbon staple, and a biaryl aromatic staple has been studied extensively in this communication using allatom molecular dynamics simulation accumulating 27 *µ*s simulation time. The free peptides have been simulated in the aqueous solution, at a higher temperature (330K) and in the presence of 8M urea to check their efficiency in restricting conformational flexibility. The p53 with the aliphatic staple is conformationally more restricted than wild-type p53 and for p53 with the aromatic staple, the flexibility is even less than that. The addition of stapling agents makes the peptides resistant to thermal and chemical denaturation also. Though the addition of stapling agent increases the hydrophobicity of the peptides, the enthalpy values do not improve. Instead the preferential binding of mdm2 with stapled peptides compared to the wild-type p53 is guided by lowering of entropic penalty which is proportional to the restriction in the conformational flexibility of the peptides. To the best of our knowledge, this is the first study of this kind where extensive quantification of the thermodynamics of p53-mdm2 binding in the presence of two stapling agents and the conformational sampling of the peptides in different environments have been combined to get a comprehensive understanding of the relation between conformational rigidification and binding efficacy. The finding from this investigation can be adopted further to design more efficient stapling agents with enhanced binding efficacy.

## Methods and Materials

### Modeling

The 13-residue stretch of human tumor suppressor p53 (sequence:ETFSDLWKLLPEN) which binds with mdm2, is considered as the system of interest. This 13-residue stretch has been used in several studies and has shown to recapitulate several aspects of the p53-mdm2 interactions.Two stapling agents were used in this investigation, one is an aliphatic hydrocarbon chain and another is a biaryl linker (Figure 1).

**Figure 1:**
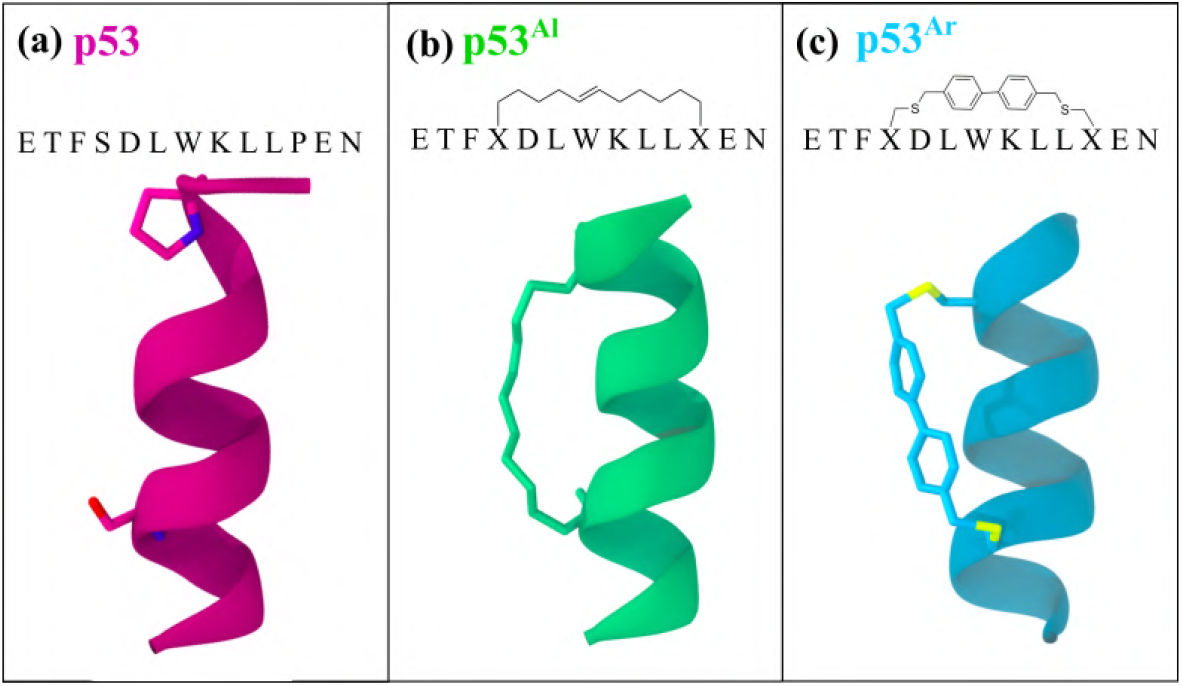
Sequences and structures of the three peptides - a) p53, b) p53^Al^, and c) p53^Ar^. The stapling positions in p53^Al^ and p53^Ar^ have been marked with X in the sequence. The peptide structures are shown in ribbon representation and the residues S4 and P11 for p53 and the stapling agents for p53^Al^ and p53^Ar^ are shown in stick representation.

The structure of wild-type p53 is taken from its crystal structure with mdm2 (Pbd Id: 1YCR).^37^ The p53 with aliphatic stapling agent attached to residues S4 and P11 is obtained from crystal structure 3V3B^29^ and is labeled as p53^Al^. The other stapled peptide is modeled based on p53^Al^ where the aliphatic stapling agent is replaced by a biaryl system connected to positions 4 and 11 via sulfur linkage. The motivation for the design is to consider a stapling agent that is different from an aliphatic staple in terms of stiffness and bulkiness. This stapling agent is termed as aromatic staple and it is termed as p53^Ar^ (Figure 1).The N- and C-terminal of the peptides were capped using Acetyl and amide group respectively to avoid interaction of the terminal charges with the rest of the system. The same capping moieties have been used in previous studies as well.^38,39^

The detailed system compositions and simulation environments have been listed in Table 1. The system identifiers defined in the table will be followed throughout the rest of the manuscript. Several simulations are performed where the wild type p53 and its stapled variations were considered in free state in different environments and also in complex with mdm2. Equilibrium molecular dynamics simulations often sample conformations that are around the global minima and thus are restricted in terms of sampling other regions of the conformational energy landscape. To overcome this and to sample conformations which differ from the native helical peptide, two unfolding inducing conditions were used - (i) Simulation at a higher temperature 330K and (ii) Simulation in the presence of 8M urea which is known to be standard and used for studying denaturation of several proteins^40,41^ and also for shorter peptides like the 13-residue stretch of p53.^42,43^ All the three peptides were simulated in these two conditions. The high temperature simulation was carried out at a slightly higher temperature than the melting temperature of p53 which is reported to be around 320K.^44,45^ To model the binding of the peptides with mdm2, three complex structures are generated - mdm2+p53, mdm2+p53^Al^ and mdm2+p53^Ar^. The first two structures are modeled according to their crystal structure 1YCR and 3V3B respectively. The mdm2+p53^Ar^ is modeled by replacing p53^Al^ with p53^Ar^ in mdm2+p53^Al^ and removing the small amount steric overlap by simple energy minimization. Simulation details of the systems are provided in Table 1.

**Table 1:**
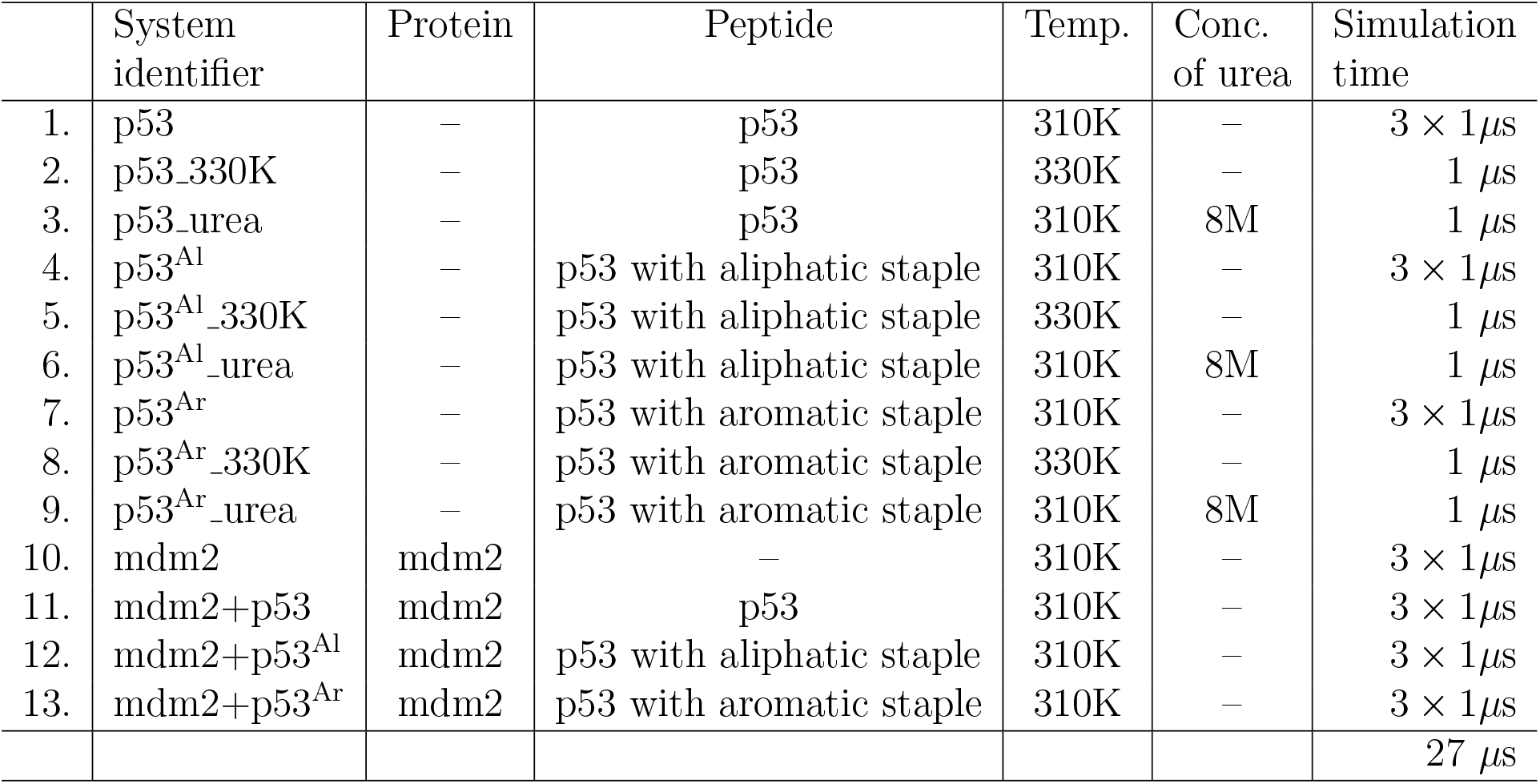
Details of the system compositions, simulation environments, and simulation times for different systems.

|  | System identifier | Protein | Peptide | Temp. | Conc. of urea | Simulation time |
| --- | --- | --- | --- | --- | --- | --- |
| 1. | p53 | — | p53 | 310K | — | $3 \times 1\mu s$ |
| 2. | p53_330K | — | p53 | 330K | — | $1\mu s$ |
| 3. | p53_urea | — | p53 | 310K | 8M | $1\mu s$ |
| 4. | p53<sup>Al</sup> | — | p53 with aliphatic staple | 310K | — | $3 \times 1\mu s$ |
| 5. | p53<sup>Al</sup>_330K | — | p53 with aliphatic staple | 330K | — | $1\mu s$ |
| 6. | p53<sup>Al</sup>_urea | — | p53 with aliphatic staple | 310K | 8M | $1\mu s$ |
| 7. | p53<sup>Ar</sup> | — | p53 with aromatic staple | 310K | — | $3 \times 1\mu s$ |
| 8. | p53<sup>Ar</sup>_330K | — | p53 with aromatic staple | 330K | — | $1\mu s$ |
| 9. | p53<sup>Ar</sup>_urea | — | p53 with aromatic staple | 310K | 8M | $1\mu s$ |
| 10. | mdm2 | mdm2 | — | 310K | — | $3 \times 1\mu s$ |
| 11. | mdm2+p53 | mdm2 | p53 | 310K | — | $3 \times 1\mu s$ |
| 12. | mdm2+p53<sup>Al</sup> | mdm2 | p53 with aliphatic staple | 310K | — | $3 \times 1\mu s$ |
| 13. | mdm2+p53<sup>Ar</sup> | mdm2 | p53 with aromatic staple | 310K | — | $3 \times 1\mu s$ |
| | | | | | | $27\mu s$ |

### Simulation details

CHARMM36 parameters^46^ for protein is used to model the peptides and the protein-peptide complexes except for the stapling agents. The stapling agents are modeled using a combination of CHARMM parameter for proteins and small molecules (CGENFF).^47^ For p53^Al^, first the side chains of S4 and P11 are replaced by that of lysine. Then the *ϵ* −NH_2_ groups are replaced by an aliphatic chain of suitable length and the end of the two side chains are joined. The carbon atoms up to *δ* carbon are parameterized using CHARMM36 protein force-field whereas the parameter for the rest is from CGENFF of CHARMM. CHARMM force field has been successfully used earlier for studying mdm2-p53 interaction.^48^ This strategy has been successfully used to model stapled peptide in several studies.^20,49^ Each of the systems is solvated in a waterbox made of TIP3P water^50^ where the size of the waterbox was determined by maintaining a distance of at least 1 nm between the water box edge and protein/peptide atoms. For the removal of initial steric clashes, a 5000-steps energy minimization was performed for each system using the steepest descent method.^51^ Subsequently, a 500 ps equilibration in the NVT ensemble was performed to equilibrate each system at 310K to avoid void formation in the box followed by a 5 ns equilibration at isothermal-isobaric (NPT) ensemble to attain a steady pressure of 1 atm. The temperature was kept constant at 310 K by applying the V–rescale thermostat^52^ and the pressure was maintained to be at 1 atm using Parrinello-Rahman barostat^53^ with a pressure relaxation time of 2 ps, used for the attainment of desired pressure for all simulations. The production runs for 1000ns with a time step of 2 fs is performed. All the simulations are performed in GRO-MACS.^54^ Short-range Lennard–Jones interactions are calculated using the minimum image convention.^55^ For estimating non-bonding interactions including electrostatic as well as van der Waals interactions, a spherical cut-off distance 1 nm is chosen. Periodic boundary conditions have been used in all three directions to remove edge effects. SHAKE algorithm^56^ is applied to constrain bonds involving the hydrogen atoms of the water molecules. Long-range electrostatic interactions are calculated using the particle mesh Ewald (PME) method.^57^ The frames in the trajectory are saved at a frequency of 2 ps for analyses. To extract different structural properties, in-built modules of GROMACS,^54^ VMD^58^ and some in-house scripts were used.

### Binding energy calculation

The standard free energy of binding (ΔG) is calculated using the MM-GBSA (Molecular Mechanics with Generalized Born and Surface Area) protocol.^59,60^ In practice, the simulations of the free and the complexed states are performed using explicit water to get the conformational microstates of the solute, and then the free energy of solvation of these conformations are estimated using an implicit environment after removal of explicit water molecules. The total free energy (G) is calculated using the following equation –

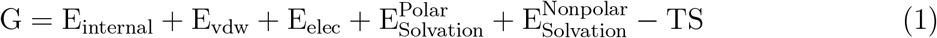

The first three terms represent the components of energy arising from the motion within the molecule (e.g., bond vibration, angle bending, dihedral rotation, etc.), van der Waals interactions, and electrostatic interactions, respectively. The fourth and fifth terms represent the solvation free energy calculated considering implicit environments in two separate parts, i.e., polar and non-polar components. The last term is solute entropy, where T represents the absolute temperature and S is the entropy of the solute, whereas the solvation entropy is already incorporated in solvation energy terms.^59^ The sum of the first five terms is considered as enthalpy, H although it includes the solvation entropy term. The values are averaged over the total simulation time and the free energy change is quantified as -

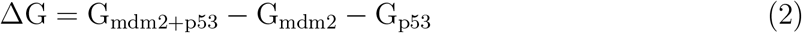

Different components of binding free energy are averaged over the last 600ns (400ns-1000ns) simulation of each system. Three independent simulations of all the systems are used for the calculation to ensure statistically significant results. Instead of using a single trajectory approach where the conformations of the components of the complex (e.g. p53 and mdm2) are derived from a single simulation of the complex (i.e. p53+mdm2)^61^ we have used a three-trajectory approach where both the components of the complex were simulated separately. This approach is proven to be more accurate in predicting binding energy.^59,62^ The values calculated for the three simulations are summarized in Supporting Information Tables (Table S1-S10) and an average of the three simulations is provided in Table 2. The implicit solvent model GBSW^63^ is used for the calculation and the dielectric constants values of the solvent and the protein interior are 80 and 1 respectively

**Table 2:** Free energies of mdm2-p53 binding. All energy values are in kcal/mol. The error bar associated with the average values for p53s, mdm2, and mdm2+p53s are provided in tables S1 to table S10)

| Energy components | p53 | p53<sup>Al</sup> | p53<sup>Ar</sup> |
| --- | --- | --- | --- |
| $\Delta E_{\text{Elec}}$ | -176.14 | -138.69 | -180.89 |
| $\Delta E_{\text{Vdw}}$ | -71.09 | -69.14 | -67.07 |
| $\Delta E_{\text{Internal}}$ | 12.46 | 13.07 | 9.04 |
| $\Delta E_{\text{Solv(polar)}}$ | 176.84 | 142.52 | 188.20 |
| $\Delta E_{\text{Solv(non-polar)}}$ | -4.41 | -4.29 | -3.93 |
| $\Delta E_{\text{Elec+Solv(polar)}}$ | 0.70 | 3.83 | 7.32 |
| $\Delta H$ | -62.34 | -56.53 | -54.65 |
| $-T\Delta S$ | 53.88 | 26.41 | 15.95 |
| $\Delta G$ | -8.46 ±0.69 | -30.12 ±2.97 | -38.70 ±2.46 |

## Results

### Stapling agent reduces conformation flexibility

The conformations of the 13-residue p53 peptide in the presence and the absence of the stapling agents obtained from the micro-second long simulation will provide an idea of the effect of the stapling agent on the behavior of the free peptide. To compare the effect of stapling agents on the conformations, a conformational energy landscape is constructed considering two important structural parameters of the peptide, backbone RMSD (root mean square deviation) and R_*g*_ (radius of gyration). The probability of occurrence of the points within 0.1 nm × 0.1 nm grid is calculated from the RMSD vs R_*g*_ plots as follows -

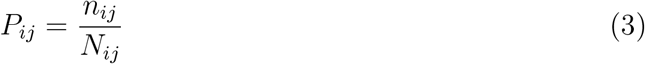

Where *n*_*ij*_ is the number of conformations within an area confined between RMSD values *i* and *i*+Δ*i* and R_*g*_ values *j* and *j*+Δ*j. N*_*ij*_ is the total number of conformations. Free energy change is then calculated using the formula –

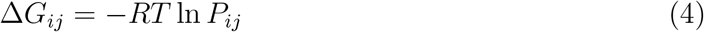

Although several other collective variables can be used to form this kind of conformational energy surface^64–66^ RMSD and R_*g*_ has been found useful in understanding the conformational transition in many cases.^67–69^ The major energy minimum points in the energy landscape have been labeled with the uppercase alphabet and the conformation of the corresponding structure has been shown in Figure 2. The plot clearly shows that the wild-type p53 has wider distribution in all the three conditions compared to p53^Al^ and p53^Ar^ systems. The distribution of the plot is restricted more along the R_*g*_ axis compared to RMSD. This can be explained by the fact that although there are deviations of the peptide conformation starting from the initial conformation in the presence of the stapling agent, the overall shape of the peptide is maintained. There are multiple minima present as separate islands on the free energy landscape of the p53 (Figure 2a) system whereas in the case of the stapled systems only one or two very closely spaced minima are present. Although in the case of both p53^Al^ and p53^Ar^ the energy landscapes look similar i.e. small energy surface distributed around one or two closely spaced minima, there is a crucial difference in their appearance in p53^Al^, p53^Ar^ and 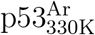 from the rest. If we considered a diagonal connecting the two most populated energy minima, along that line both RMSD and R_*g*_ increases simultaneously. Whereas, in case of p53^Al^, p53^Ar^ and 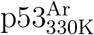 the diagonal is opposite in direction i.e. conformation with higher R_*g*_ is associated with lower RMSD. Thus the addition of a stapling agent can successfully reduce the structural changes quantified in terms of both RMSD and R_*g*_. To find the detail in the residue level fluctuation, the root mean square fluctuation (RMSF) is calculated and the values for different systems are compared in Figure 2(j)-(l). It shows that the restriction of the residue level fluctuation is not limited to the residues between the stapling positions (i.e. S4 and P11) rather it propagates all over the peptide.

**Figure 2:**
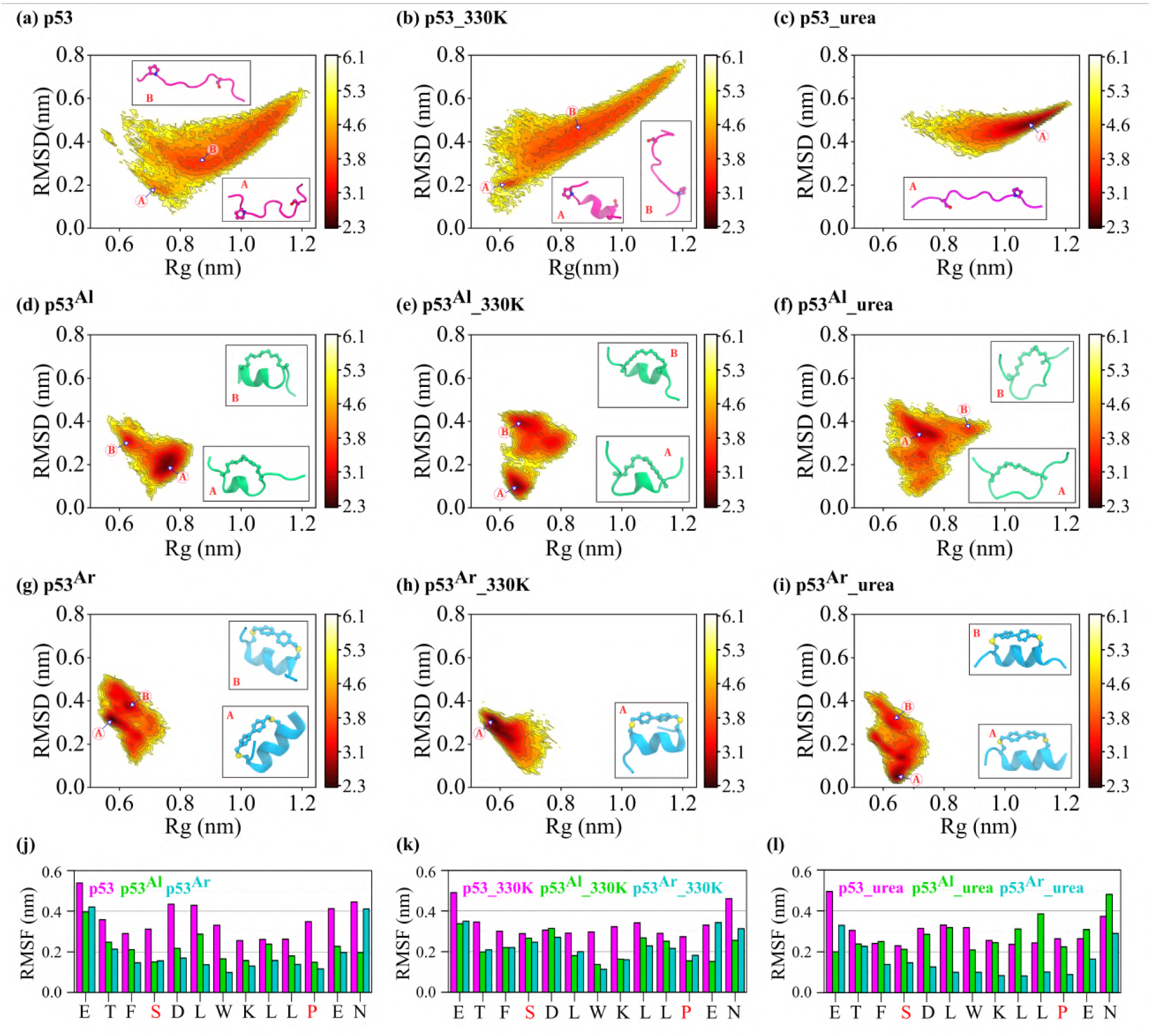
2D Free Energy Surface (FES) obtained from RMSD (Root mean square deviation)-vs-R_*g*_ (Radius of gyration) plot for the p53 peptide and the p53 peptide with aliphatic and aromatic staples in different conditions (i.e. in the aqueous medium, at high temperature, and in the presence of urea) showing conformational restriction in the presence of stapling agents. The dark red-to-yellow-to-white color scale represents increasing conformational free energy in kcal/mol. The major energy basins in the energy profiles are marked with upper case alphabets (A, and B) and the conformations of the peptide corresponding to the energy basins are shown inside the plot within rectangular boxes. The root mean square fluctuations (RMSF) of each residue of the peptides have been plotted together for the three conditions - (j) in the aqueous medium (k) at high temperature, and (l) in the presence of urea in separate graphs. The residues replaced by the stapling agent i.e. S4 and P11 are colored red.

The peptide binds to mdm2 to occupy its binding pocket in a conformation which is a helix. The helical conformation is crucial for the orientation of the binding pocket anchoring residues of p53 e.g F3, W7, and L10. Thus it will be interesting to look into the helical propensity of the peptide in the presence of thermal and chemical denaturing agents and how that is altered when a stapling agent is introduced.

### Stapling enhances backbone rigidity to restrict unfolding

The helical propensity of the peptide is quantified as fraction helicity. First the number of six residue *alpha*−helical stretch (S) in the peptide is calculated using PLUMED^70^ according to the following formulae as introduced by Pietrucci *et al*.^71^ -

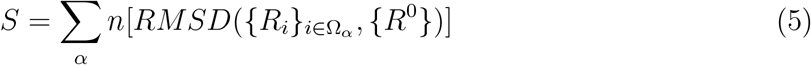

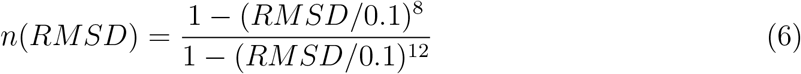

where {*R*_*i*_}_*i*∈Ω*α*_ are the coordinates of the set Ω_*α*_ describing a six residue protein-stretch and {*R*^0^} are the corresponding coordinates of an ideal *α*-helix. The number *S* is then divided by the maximum number of possible six residue *α*-helical stretches in the peptide, *S*_*max*_ to get the helical fraction (*f*_*H*_)

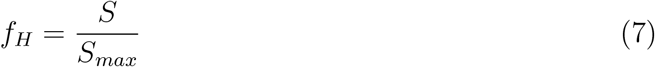

The time evolution of *f*_*H*_ is calculated for the peptides in different system and presented in Figure 3. In Figure 3a, the helical fractions of p53, p53^Al^, and p53^Ar^ in aqueous solution were plotted together.

**Figure 3:**
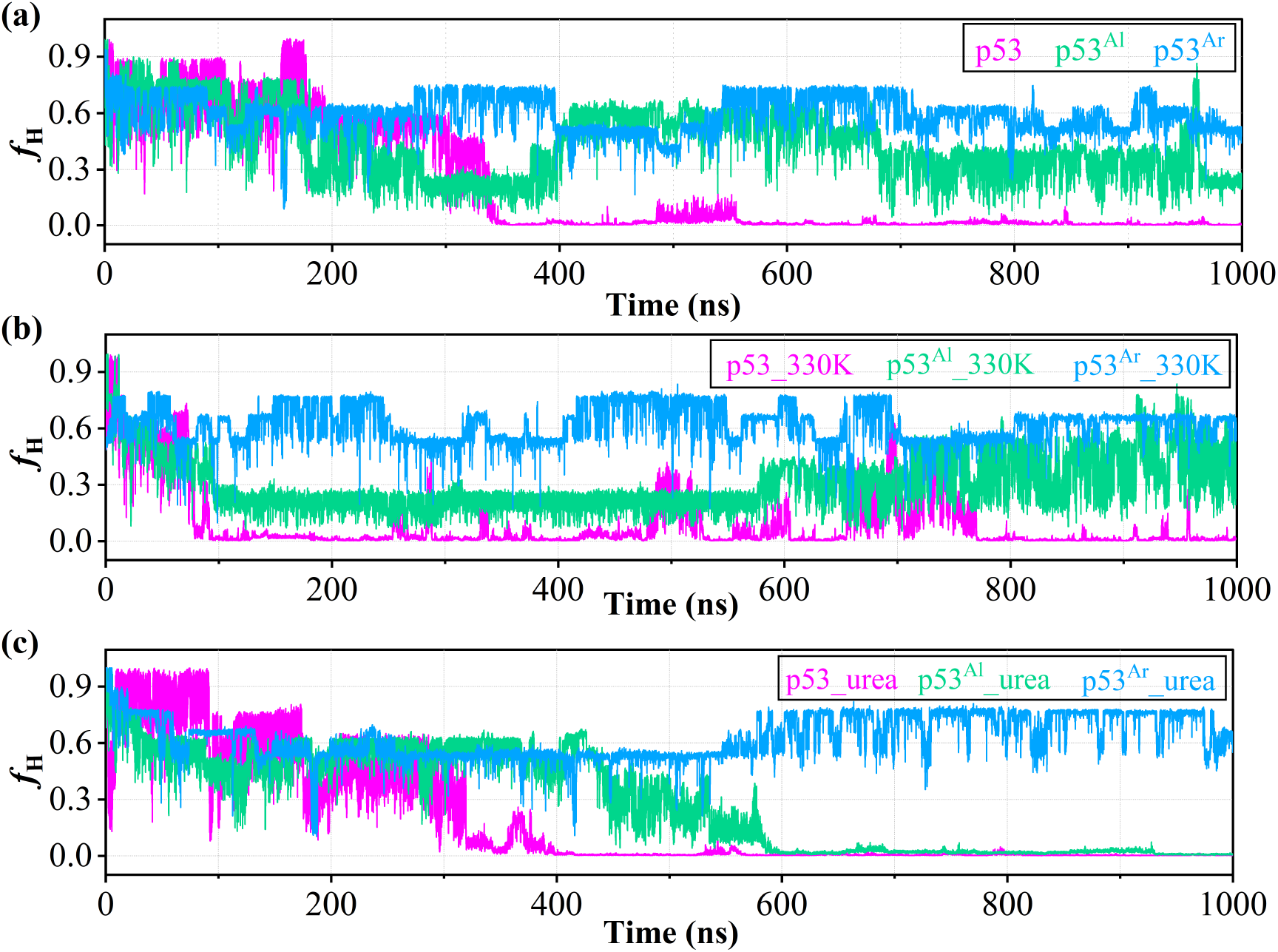
Time evolution of helical fraction (*f*_H_) of p53, p53^Al^, and p53^Ar^ in a) aqueous solution, b) at high temperature and c) in the presence of urea. The *f*_H_ is calculated according to equation 7. The p53 loses helicity in all the conditions (magenta lines), p53^Al^ maintains the helicity in aqueous environment and at high temperature but loses helicity after 600 ns in the presence of urea (green lines) whereas p53^Ar^ remains in helical conformation in all conditions (cyan lines).

It can be seen that the helical fraction of p53 gradually decreases to zero at 350ns and rest of the simulation, the peptide was in a coiled conformation. Upon addition of stapling agents the helical fraction was maintained and it was more effective for p53^Ar^ compared to p53^Al^. The helicity measured using experimental techniques by Zondlo *et al*.^72^ and Xiong *et al*.^73^ show the wildtype p53 to have considerable extent of helical structure which is in agreement with the initial part of our simulation but not with the unfolded ensemble. On the other hand, Bernal *et al*.^18^ reported the helicity of the wild type p53 to be 11% which corresponds to an unfolded peptide and it matches well with the conformations from our simulated ensemble. An introduction of an aliphatic staple has improved the helicity to 60% in the same study and the enhancement in helicity for p53^Al^ in the present study agrees with that result.

Additionally, the calculated helicity from our simulation is in agreement with previous computational studies by Guo *et al*.^74^ where they found an improvement in the helicity (from 2% to 38%) upon addition of aliphatic stapling agent to p53 and Sim *et al*.^75^ where helicity was improved significantly upon addition of the stapling agent. To check the effectiveness of the stapling agents in maintaining the helicity, two external unfolding agents were employed to facilitate the unfolding of the peptide. The peptides were simulated at 330 K to model thermal denaturation and to model chemical denaturation, the peptides were simulated in an 8M urea solution. At 330K the p53 unfolds at an earlier time (100ns). The correlation between the change in the helical propensity and in the global conformational change quantified using RMSD and R_*g*_ is verified using RMSD-vs-helicity and R_*g*_-vs-helicity plots (Figure S1 and S2). A careful observation of the plots reveals that the energy minima for p53 in all the three conditions are associated with very lower value of helicity and higher values of Rg representing a higher population of the unfolded conformation. On the other hand, for the p53^Al^ system the highly populated with conformations having moderate value of both helicity and Rg. In case of p53^Ar^, the most populated conformations have lower Rg and higher helicity compare to both p53 and p53^Al^. The addition of aliphatic and aromatic staples can reduce the extent of unfolding where aromatic stapling agent is found to be more effective than the aliphatic one. To check the exposure of the peptide backbone to the solvent, the radial distribution functions of water and urea molecules around backbone atoms and the number of peptide-water and peptide-urea hydrogen bonds were calculated and shown in Figure 4. In all the cases the wild type p53 systems show maximum abundance of solvent around their backbones whereas it decreases as the stapling agents are added. In aqueous solution and at 330K, both p53^Al^ and p53^Ar^ have distribution of water significantly below that of the p53 (Figure 4a-i and Figure 4b-i). In the presence of 8M urea, the distribution of water and urea for p53^Al^ is close to that of p53 while p53^Ar^ shows a lower distribution of water and urea (Figure 4c-i and Figure 4d-i). The p53 forms maximum number of hydrogen bonds with water or urea in all the systems which reduces upon addition of the stapling agents (Figure 4a-ii-4d-ii). This differences in RDF and hydrogen bond number signify that the p53^Ar^ system is more resistant to both thermal and chemical denaturation whereas the p53^Al^ is less effective to reduce the extent of unfolding in the presence of urea.

**Figure 4:**
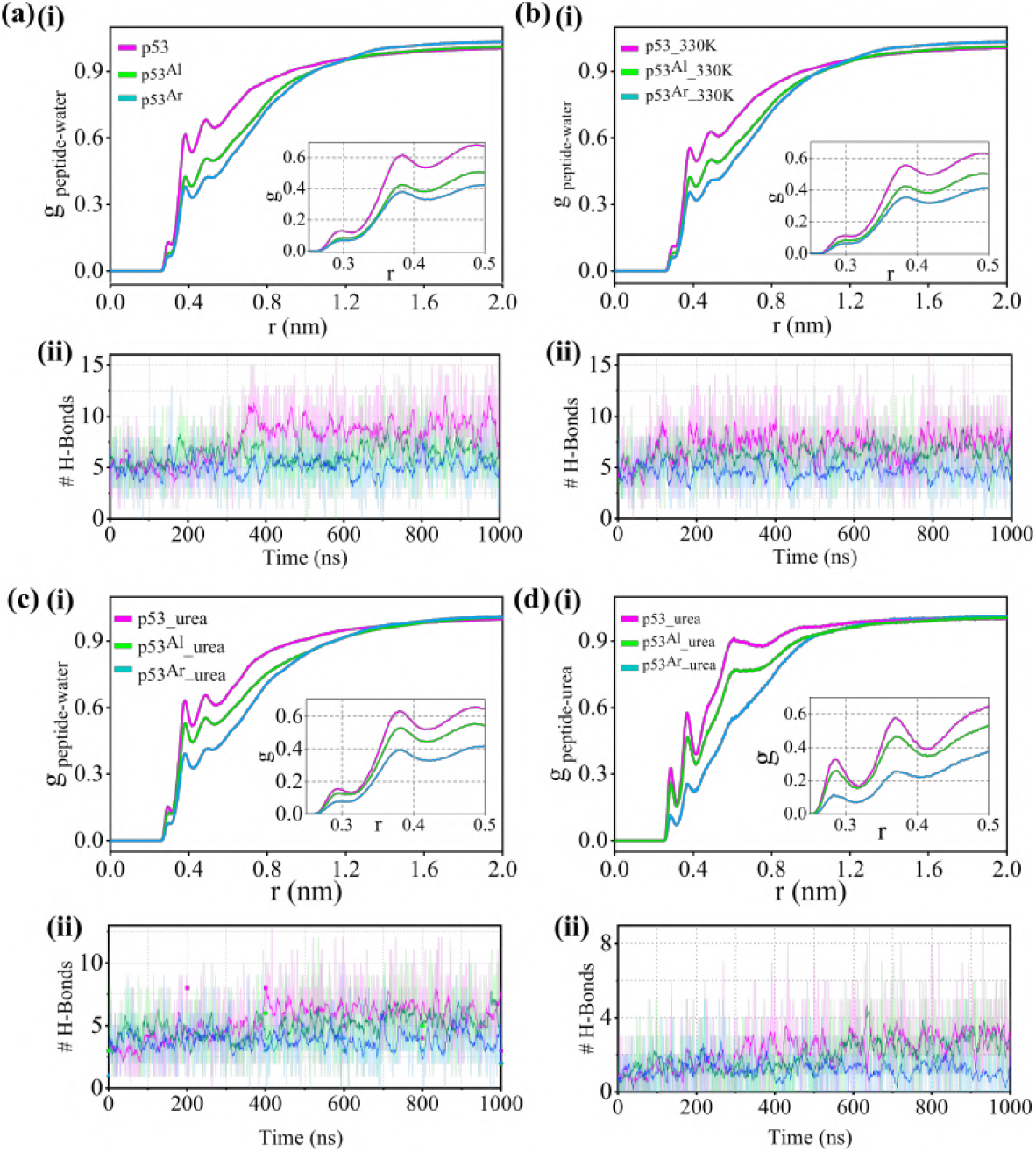
Radial distribution function (rdf) of water around peptide for different systems in a-i) aqueous solution, b-i) at 330K, and c-i) in the presence of urea. The rdf of urea around protein for all three systems is shown d-i). The rdf is calculated for the center of mass of water or urea molecules around the backbone atoms of protein (i.e. C, N, and CA). In all cases, p53 has highest value of rdf because of the higher accessibility of the backbone atoms due to unfolding (magenta lines). Rdfs for p53^Al^ and p53^Ar^ appear below that of p53 ensuring the lower solvent exposure of backbone atoms and hence maintaining the helical conformation (green and cyan lines respectively). The number of hydrogen bonds between water molecules and backbone atoms of different peptides throughout the simulation in a-ii) aqueous solution, b-ii) at 330K, and c-ii) in the presence of urea are shown. The hydrogen bonds between peptide backbone and urea molecules for three systems during the simulation are presented in d-ii). In each case, addition of stapling agent reduces the number of hydrogen bonds representing more folded conformation.

Overall the structural changes of p53 and its stapled variants in water and in the presence of thermal and chemical denaturant show that there is a significant reduction of the conformational flexibility when the S4 and P11 side chains were modified and joined together like a stapled residue. This restriction in the conformations and modification of the amino acid side chain of p53 will influence the binding to its partner mdm2. Although the effect can influence the binding in many ways it could be categorized into two major types - (i) The interaction of the polar S4 and P11 with mdm2 binding pocket residue will be compromised and instead there will be additional interaction of hydrophobic stapling agent with the binding pocket. A resultant of these will be reflected in the protein-peptide interaction. (ii) The energetic cost to bring the peptide from a free state to a mdm2-bound state will be different when the conformation in the free state itself is modified. To understand these in detail, the free enegry of mdm2-p53 binding has been calculated using MM-GBSA protocol. Although there are other methods for binding energy calculation,^36,76^ they are computationally more expensive and are not extensively used to study protein-peptide binidng. Whereas, MM-GBSA is a well standardized protocol for calculating binding free energy and it is widely used.

### Entropy determines the binding affinity

The binding free energy was estimated considering one microsecond long simulation of mdm2+p53, mdm2, and p53 systems. The average binding free energy was quantified considering both enthalpy and entropy. The enthalpy was calculated using the MMGBSA protocol, a formalism used to find the solvation energy using a continuum model for solvent. The conformational entropy is estimated considering quasi harmonic approximation on the ensemble of conformations. These values are further averaged over the three simulations of the protein+peptide systems. The results for the three systems are summarized in Table 2. The values of separate simulations are summarized in the supporting information Tables. The MMGBSA formalism is successfully used for the calculation of binding free energies of protein-small molecule and protein-peptide binding and shows reasonable agreement with the experimentally determined binding energies.^77^ There are also successful applications of this method in estimating the binding energies of small molecules,^78^ peptides^79^ and stapled peptides^32^ with mdm2. At this point it is important to note that the relative difference in the calculated binding energies of the systems under investigation is more relevant than the absolute binding free energy values.

The components of binding energy show that the overall binding energies p53^Al^ and p53^Ar^ are more favorable than p53. Interestingly, the enthalpy and entropy values follow the opposite order. Despite the addition of the stapling agent the enthalpy is found to be reduced for the stapled systems. The analyses of individual energy components of enthalpy show that the lowering of enthalpy originates mainly from two quantities, ΔE_Vdw_ and ΔE_Elec+Solv(polar)_ representing the change in van der Waals energy and total polar interaction energy respectively. There is a loss of ~2 and ~4 kcal/mol in the van der Waals interaction energy for p53^Al^ and p53^Ar^ respectively when the stapling agents are added. The ΔE_Elec+Solv(polar)_ is a sum of ΔE_Elec_ and and ΔE_Solv(polar)_. The first term is the change in electrostatic interaction upon complexation. The second term is the change in polar solvation energy between the protein-peptide complex in the aqueous medium and the protein and the peptide separately. The values of these two terms are usually complementary to each other i.e. if there is a gain in the interaction energy upon binding it is usually associated with a loss in the polar solvation energy because the protein-solvent polar interactions for the residues forming the binding interface are replaced by the protein-protein electrostatic interactions. Here the electrostatic interaction is reduced in the stapled peptides and the loss in polar solvation (i.e. positive ΔE_Solv(polar)_) decreases (mainly for p53^Al^). A combination of these results in an overall unfavorable change in polar interaction for the stapled peptides compared to the wild-type p53. In contrary to enthalpy, the TΔS follows the reverse order. The value of TΔS itself is negative here i.e. going from the ‘free’ state to the ‘mdm2-bound’ state the peptide loses entropy. Interestingly the amount of entropy loss has been significantly reduced when the stapling agents were added. This reduction in entropic penalty has significantly contributed to the overall free energy. Overall the binding of stapled p53 peptides is thermodynamically more favorable than the wild-type peptide.

The experimentally determined binding energy values for p53 with mdm2 lie in the range of −6.8 to −9.1 kcal/mol^80–83^ and the value obtained from the present study (−8.46 kcal/mol) are in good agreement with that. On the other hand, binding energy values obtained from some previous computational studies are −6.4 kcal/mol^32^, −7.4 kcal/mol^84^ and −12.9 kcal/mol.^85^ The difference between the predicted and experimental binding energy values arises from a number of reasons such as force field parameters, binding energy calculation method, simulation length etc. For example, Zhou *et al*.^84^ and Paul *et al*.^85^ have used extensively long simulations and other statistical means. Whereas, we have used relatively simpler and computationally less expensive techniques yet the binding energy value is in good agreement with the experiment. However, the deviation of computationally predicted values from experimental values by an amount of 1-2 kcal/mol has also been argued to be in the limit of force field inaccuracies.^85,86^ Moreover the effectiveness of the stapling agents to enhance the binding affinity, found here is in agreement with experiments and previous computational studies.

Since both the free state of p53 and its mdm2-bound conformation have roles to play in determining the binding efficiency, a comparison of the important features of mdm2 binding in the two states will be insightful to rationalize the binding free energy values. In the next section, two such key features will be discussed in detail.

### Stapling reduces the cost of conformational transition upon binding with mdm2

The crystal structures of p53-mdm2 complexes show that the binding of p53 with mdm2 is guided by anchoring of three hydrophobic residues into the binding pocket which are F3, W7, and L10 and the p53 peptide fit into the binding pocket with a helical conformation. These two important features i.e. the helicity of the peptide and the orientation of the three residues were monitored in the free and mdm2-bound state of p53. A comparison of helicity of the free and mdm2-bound conformation is presented in Figure 5. For p53, the helical fraction in the mdm2-bound state is maintained around 0.5 whereas the free state looses its helicity after 350ns. Thus the transition from free to mdm2-bound state will require large amount of energy for unfolded → folded transition. While considering the aliphatic stapled system, although there are fluctuations in the helical fraction there are significant overlaps between them and the average *f*_*H*_ values remain in the same range. Interestingly p53^Ar^ and mdm2+p53^Ar^ have higher *f*_*H*_ value throughout the simulation with good overlap between them. This reduces the energetic cost required to change the peptide conformation prior to binding with mdm2 making the process more feasible compared to that for p53. A qualitative estimation of this energy is shown in figure S3. The Rg and helicity values corresponding to the mdm2-bound conformations of the peptides overlap on the Rg-vs-helicity free energy surface (FES) of the free p53 systems. It shows that the conformations corresponding to the mdm2-bound p53 is far away from the energy minimum of the FES (Figure S3a). Whereas the same plot for p53^Al^ and p53^Ar^ shows a significant overlap of mdm2-bound conformation with the energy minimum where the extent of overlap is higher for p53^Ar^ (Figure S3c) than p53^Al^ (Figure S3b). A comparison of structural rigidity of the peptide in the binding pocket of the protein can be well understood from the visualization of the simulation of mdm2+p53 (Movie S1), mdm2+p53^Al^ (Movie S2), and mdm2+p53^Ar^ (Movie S3) respectively.

**Figure 5:**
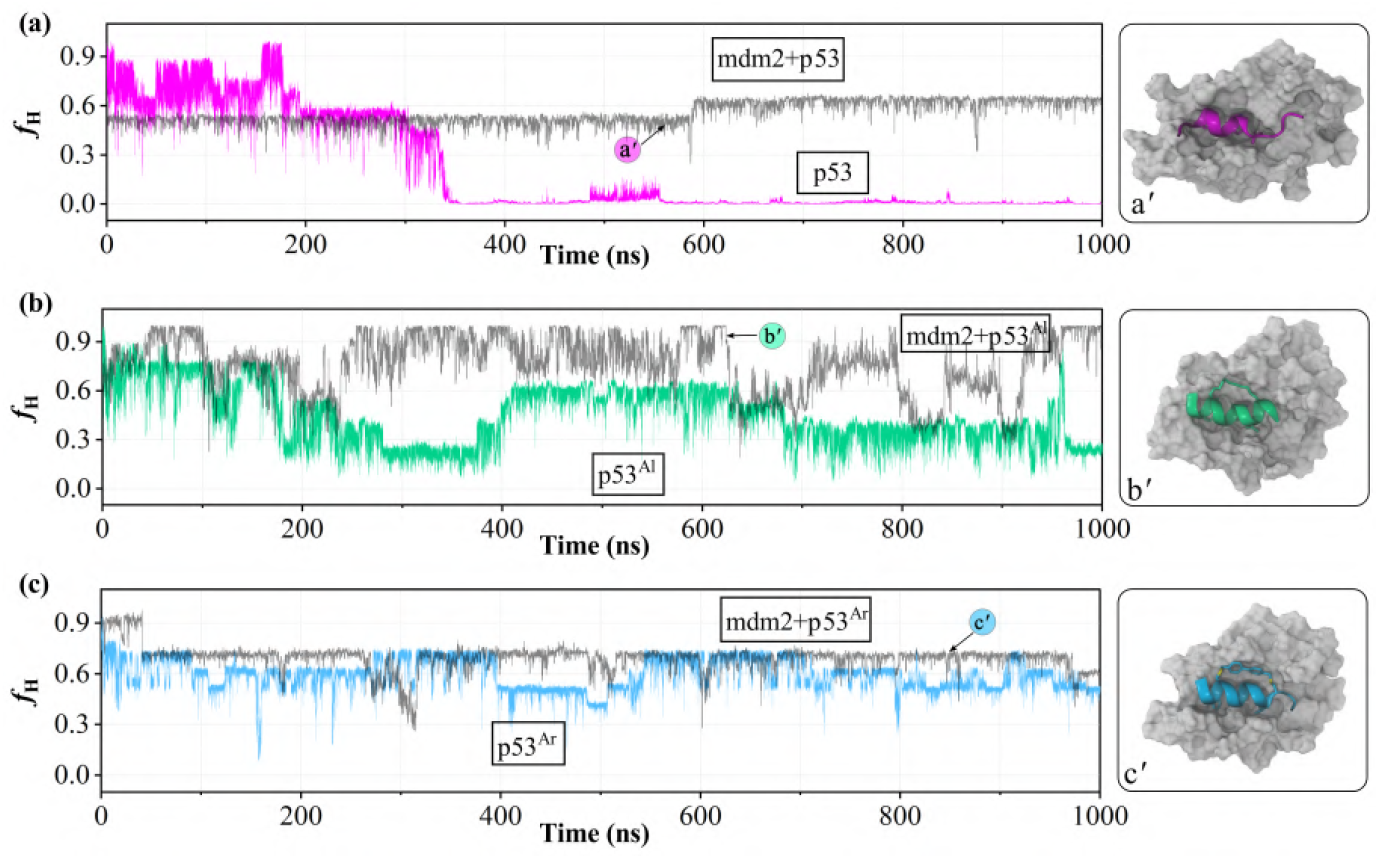
Time evolution of helical fraction (*f*_H_) of a) p53, b) p53^Al^, and c) p53^Ar^ in free and mdm2-bound state. The *f*_H_ of mdm2-bound state is shown in gray lines in each plot. The *f*_H_ is calculated according to equation 7. The helicity of p53 in free and mdm2-bound state is significantly different (a) whereas they have considerable overlap for p53^Al^ (b) and high degree of overlap for p53^Ar^ (c). A representative conformation of each of the mdm2-bound states of the three peptides is shown next to the plots. The mdm2 is shown in light gray surface whereas the p53, p53^Al^, and p53^Al^ are shown in magenta (a), green (b), and cyan (c) ribbon respectively.

Since the peptide has gained enhanced helicity inside the mdm2-binding pocket, the interaction of the peptide with the binding pocket residue may also get modified. To quantify that, the interaction energies of each residue of the peptides are calculated with mdm2-binding pocket residues and shown in Figure 6. It is evident from the plot that the interaction of the three anchoring residues (F3, W7 and L10) has not changed much upon addition of the stapling agent. Whereas, there is a significant improvement of the interaction when interaction of S4 and P11 of p53 with mdm2 is compared with the interaction of the stapling agent of p53Al and p53Ar with mdm2. Interestingly an increase in the helicity upon addition of stapling agent enhances the interaction of C-terminal residues (E12 and N13) of p53 with mdm2.

**Figure 6:**
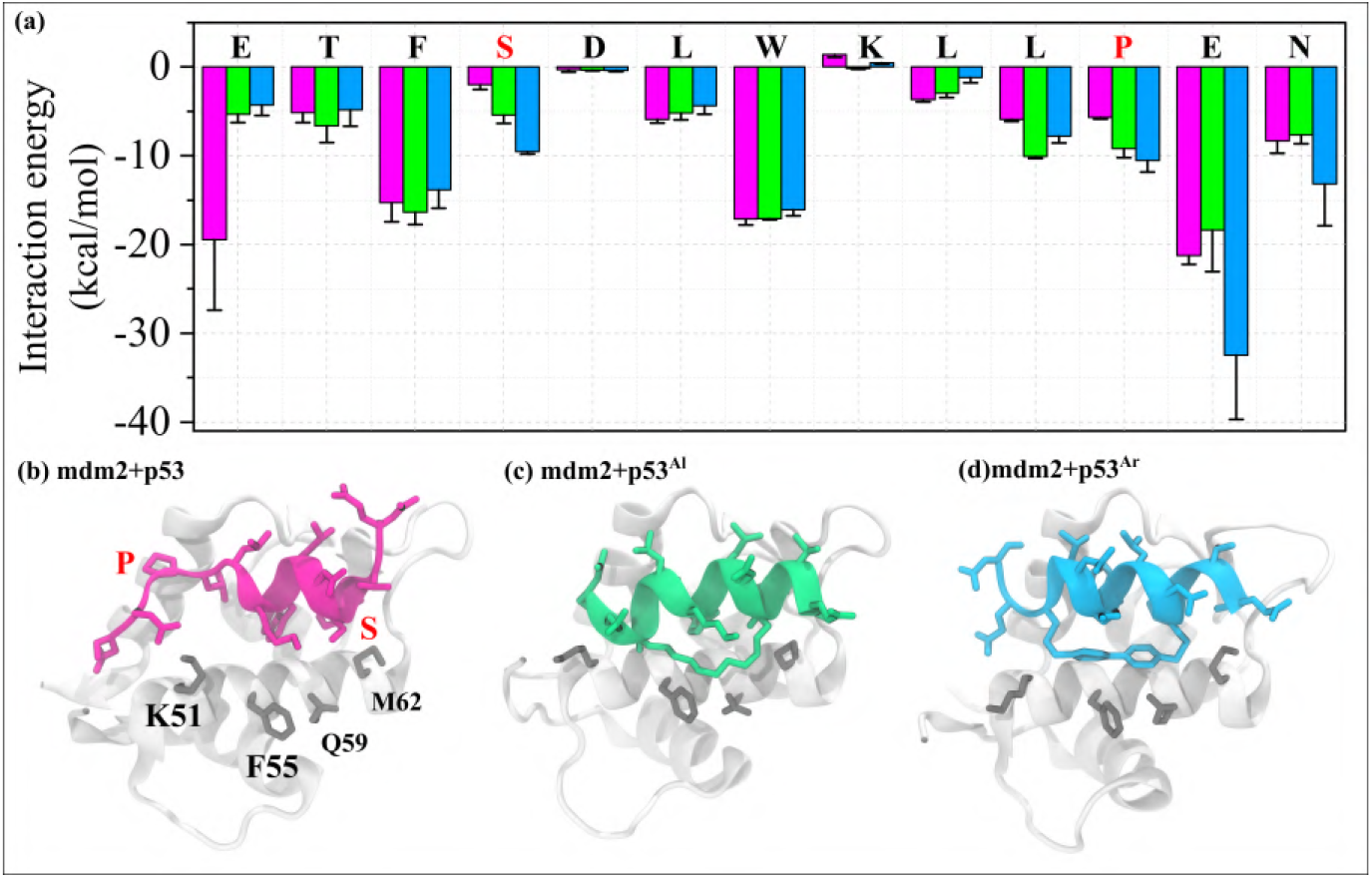
(a) Interaction energy between each amino acid of the peptide and the binding pocket residues of mdm2. The residues which are replaced by the stapling agent are shown in red. Representative structures of the complexes have been shown for (b) mdm2+p53, (c) mdm2+p53^Al^, and (d) mdm2+p53^Ar^. The mdm2 is shown in white ribbon in all of the structures and the residues directly interacting with the peptide are in gray stick.

To compare the relative orientation of the three anchoring residues, two distances among them have been monitored throughout the simulation time which are *r*_*F*3−*W*7_ and *r*_*W*7−*L*10_. A plot of *r*_*F* 3−*W* 7_-vs-*r*_*W* 7−*L*10_ is shown in Figure 7. The magenta points in Figure 7b represent the distances for the free p53 and the points encircled in black are distances calculated from the mdm2+p53 system. An overlap of these shows that the free peptide deviates by a large distance in terms of the favorable distances for complexation. As a result, the energetic cost of reorienting the residues as suitable for complexation will be energetically costly. This overlap for the p53^Al^ and p53^Ar^ systems depicts that the points for free and mdm2-bound states are quite close to each other, specifically for p53^Ar^ system. This ensures an energetically favorable transition from free to complexed state. The effect of the stapling agent in maintaining this orientation is even more evident if the free peptide simulations in different conditions are considered (Figure S4).

**Figure 7:**
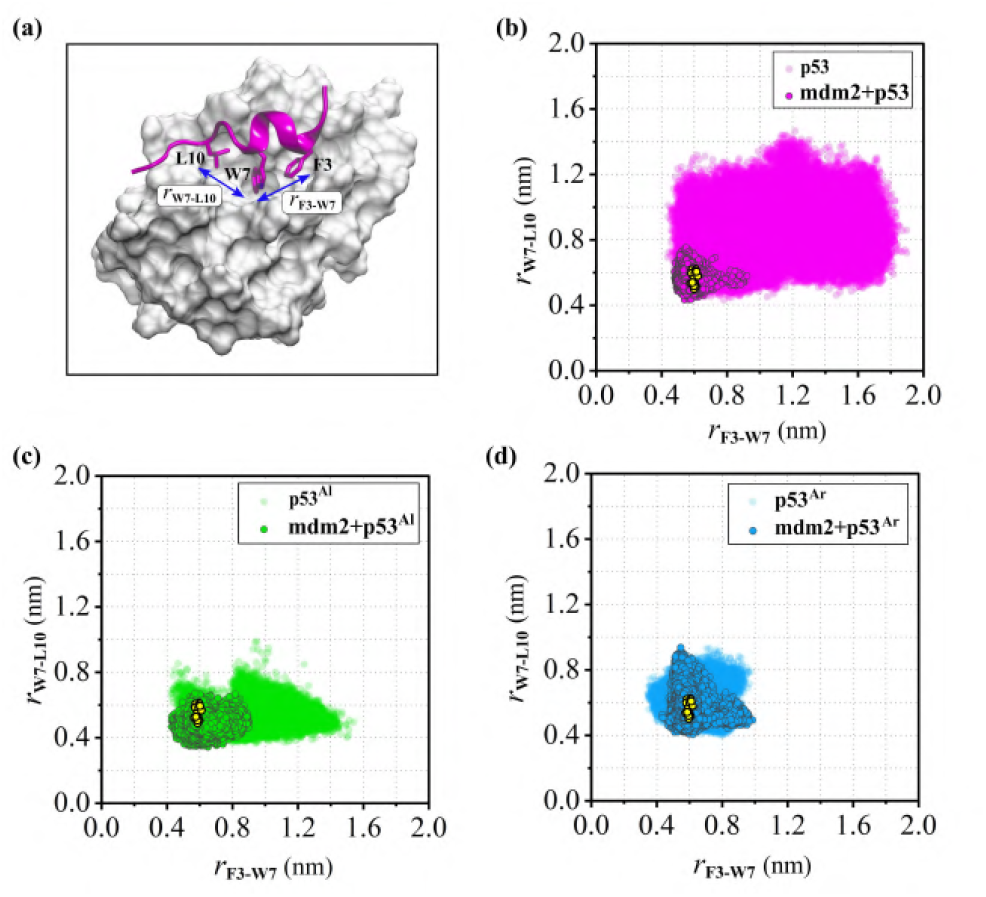
a) Anchoring of F3, W7, and L10 (magenta sticks) into the binding pocket of mdm2 (white surface). Overlap of the plots of *r*_*F*3−*W*7_-vs-*r*_*W*7−*L*10_ for the free and mdm2-bound states of b) p53, c) p53^Al^, and d) p53^Ar^. Distances for the mdm2-bound state are plotted as circle with gray circumference while those from the free state has no circumference. The free-state conformations of p53 is widely distributed and it has very small overlap with the mdm2-bound state (b) whereas for both p53^Al^ (c) and p53^Ar^ (d) the free and bound state distance distributions overlap with each other. This signifies lower costs of conformational transition upon mdm2-binding for p53^Al^ and p53^Ar^ compared to p53. For comparison of the simulated structures with the crystal structures, the same pair of distances from twenty eight crystal structures of different p53/modified p53-bound mdm2 are shown as yellow dots on the plots.

## Discussion

The conformational dynamics of wild-type p53 and its two stapled versions p53^Al^ and p53^Ar^ have been monitored in their free states in water and in their complexes with mdm2. The peptides are more flexible in the free state than when they are bound to mdm2 binding pocket and this flexibility is also influenced by the addition of stapling agents. The conformational flexibility has been measured using two properties, RMSD and R_*g*_. The plot clearly shows that change in both RMSD and R_*g*_ is significantly large and the R_*g*_-vs-RMSD plot spans over a wide area (Figure 2a(i)). This is significantly reduced when the stapling agent is added by replacing the amino acids S4 and P11 (Figure 2b(i) and Figure 2c(i)). This reduction in overall structural change is initiated by the cross-linking of S4 and P11 and a restriction in the backbone atom motion in that region can influence the helical propensity of the peptide. This is confirmed by comparing their helicities during the simulation. The p53 loses its helicity after 350ns whereas the stapled peptides maintain the helical fraction in the range 0.4 to 0.7 throughout the simulation (Figure 3a). Since the equilibrium simulation of the peptide in water has its limitation in sampling all possible conformations, two external conditions are imposed that can accelerate the sampling of conformations distant from the stable helical conformation. This will sample the unfolded conformations of the wild-type p53 and also check the endurance of the staple peptides in these adverse environments. The result shows that indeed they are capable of restricting the conformational sampling (Figure 2). In the condition of thermal denaturation both p53^Al^ and p53^Ar^ are capable of preventing the unfolding (Figure 3b) whereas in the presence of urea, there is significant loss of helicity of the p53^Al^ after 600ns (Figure 3c).

To estimate the influence of these stapling agent-induced conformational changes on the binding of p53 with mdm2, two important structural factors of p53 are checked that facilitate its binding. One of them is the helical structure of the peptide and another is the relative orientation of the three anchoring residues F3, W7, and L10. The values of these two structural parameters are compared for the free peptides and their mdm2-bound states in Figure 5 and Figure 7 respectively. The difference between the average helicities of the peptide in the mdm2-bound state and the free state is a determinant of the relative ease with which it can bind with mdm2 considering a population shift model^87^. These differences (Δ*f*_*H*_) are approximately 0.7, 0.3 and 0.1 for p53, p53^Al^, and p53^Ar^ respectively. Thus the energetic barrier for conformational switching from free to complex state follows the order - p53^Ar^ < p53^Al^ < p53. A closer look into the distances between the anchoring residues *i*.*e. r*_*F* 3−*W* 7_ and *r*_*W* 7−*L*10_ in free and complex states indicates the maximum overlap for p53^Ar^ (Figure 7c) and a significant overlap for p53^Al^ (Figure 7b) whereas for p53, the points for complex state overlap only with a small part of the free state distribution (Figure 7a). The advantages of cross-linking two amino acid side chains quantified through the above structural parameters are further verified by considering the calculation of thermodynamics of mdm2-p53 binding. The binding free energy follows the trend 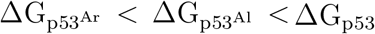. This justifies the idea that restraining the peptide conformation and inducing helical propensity using a stapling agent can enhance the binding affinity. The experimental binding affinity of the p53 and p53^Al^ are reported to be 700 nM and 55 nM respectively which follows the same order found from our calculations.^29^ The origin of this enhanced free energy of binding can be understood if the enthalpy and entropy of the binding process are considered separately. Although the addition of stapling agent results in a compact conformation of the peptides inside the binding pocket, the binding enthalpy is increased (*i*.*e*. unfavorable). The hydrophobic nature of the long stapling agent spanning over seven residues induces significant van der Waals interaction with the protein but is not sufficient to compensate for the loss of electrostatic interaction. The gain in helicity of the peptide due to the introduction of the stapling agent at the 11th position induces conformation restriction to the residues E12 and N13 also. As a result it loses some of the polar interaction with mdm2 in mdm2+p53^Al^ as well as in mdm2+p53^Ar^. This results in the increase in ΔH value (Table 2). Interestingly the change in entropy plays an important role to recover this loss. The change in conformational entropy (ΔS) is defined as -

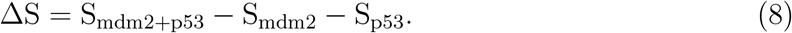

Due to the large conformational freedom of the free peptide, *S*_*p*53_ has a very high value and that results in an overall negative Δ*S*. The addition of a stapling agent has reduced the conformational flexibility of the free peptide as reflected in the R_*g*_-vs-RMSD plot and in the helicity values (Figure 2 and 3a). This reduces the free state entropy value of p53^Al^ and p53^Ar^ significantly. Since the conformational fluctuations of the complex states (the first term at the right-hand side of the equation) are almost similar for the three complexes except for the difference in some of the amino acids at the binding pocket, the major changes in ΔS arise from the entropy of the free peptide (the third term at the right-hand side of the equation). Due to the smaller values of S_p53Al_ and S_p53Ar_ the ΔS_p53Al_ and ΔS_p53Ar_ are less negative than ΔS_p53_. This decrease in the entropic penalty is large enough to overcome the increase in enthalpy and the free energy of binding decreases. This type of enthalpy and entropy compensation has been observed by previous studies as well.^88–90^ This favors the binding of stapled peptide over the wild-type p53. Thus the conformational restrictions of free peptides checked using simulations in several conditions have their influence on the entropy values. The structure of the two stapling agents and the thermodynamic parameters justify that a restriction of the peptide conformation is fruitful in enhancing the binding affinity and that the stiffness of the stapling agent is beneficial for the binding. Like most of the binding free energy calculation methods, MM-GBSA method also has limitations. In particular, MM-GBSA method sometimes fails to produce very accurate absolute ΔG values and it depends on the internal dielectric constant of the solutes.^91^ However, the method has shown remarkable accuracy in assessing the relative order of protein-peptide binding energies in different contexts.^77,92^ The understanding of the chemical nature of the stapling agent and its binding efficiency can be used as a guiding principle to design more effective therapeutics against mdm2 in the future. A successful adoption of the strategy described in this work can lead to opening of new avenues towards designing therapeutics against many other diseases.

## Supporting information

Coordinate files

## Associated Content

Supporting Information: 2D histograms of RMSD-vs-helicity and R_*g*_-vs-helicity; R_*g*_-vshelicity overlaps between free peptides and peptide-mdm2 complexes; Orientations of anchoring residues in different conditions; Detailed average energy components for all systems with standard errors; Dynamics of p53, p53^Al^, and p53^Ar^ at the binding pocket of mdm2.

## Data and Software Availability

The coordinates (.pdb files) of proteins and peptides used for the simulations of the systems listed in Table 1 are available in the Supporting Information. The sofwares, GROMACS-2018.8 (used for MD simulation), PLUMED-2.4.3 (used for trajectory analyses) VMD-1.9.3 (used for structure visualization, rendering movies and snapshots) and Inkscape-1.0.2 (used for preparing figures) are available for free. The academic version of CHARMM-42b1 is used for the preparation of systems for simulations and for analyses. Origin-2020 is used for plots through the license owned by Indian Institute of Technology Bombay.

## Acknowledgments

RC acknowledges SERB (Grant CRG/2020/000279) for financial support. AM acknowledges Indian Institute of Technology Bombay for Postdoctoral Fellowship. ARC achnowledges DST, Inspire for the fellowship. Authors acknowledge the computational facility (HPC) provided by Indian Institute of Technology Bombay. Authors thank Dr. Sandeep Choubey from Max Planck Institute for the Physics of Complex Systems and Mr. R. Kailasham for critical reading of the manuscript.

## Table of content graphics

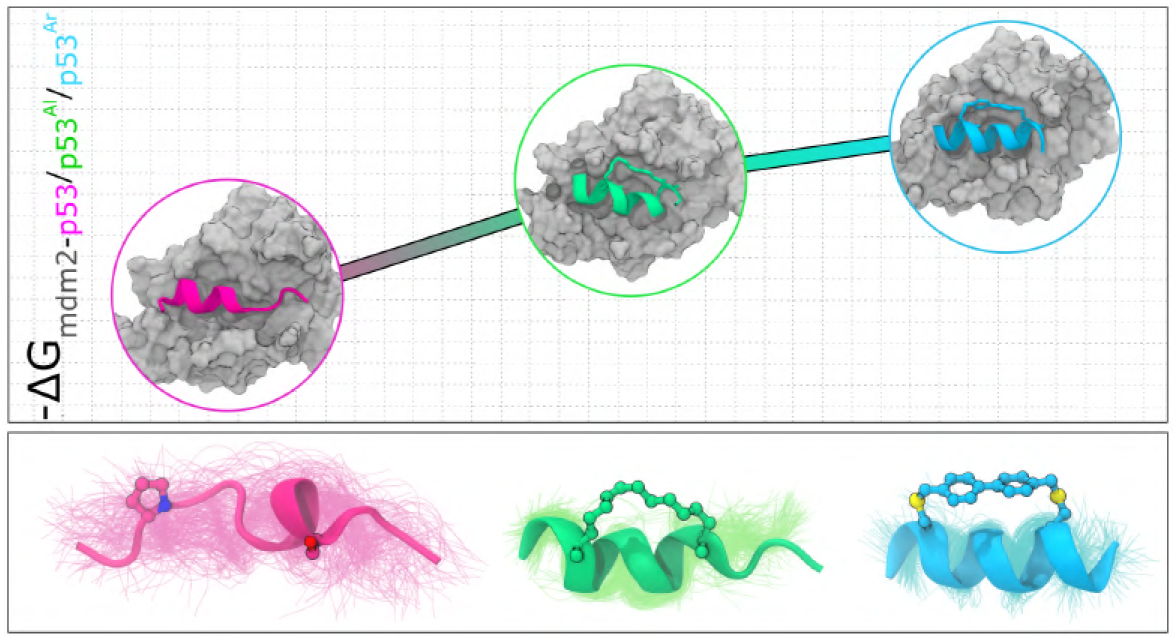

## References

(1) Cooper, G.M., Hausman, R.E. The Cell: Molecular Approach; Medicinska naklada, 2004.

(2) Ryan, D.P., Matthews, J.M. Protein–protein Interactions in Human Disease. Curr Opin Struct Biol 2005, 15, 441–446.

(3) Zinzalla, G., Thurston, D.E. Targeting Protein–protein Interactions for Therapeutic Intervention: A Challenge for the Future. Future Med. Chem. 2009, 1, 65–93.

(4) Yang, J., Hu, L. Immunomodulators Targeting the PD-1/PD-L1 Protein–protein Interaction: From Antibodies to Small Molecules. Med. Res. Rev. 2019, 39, 265–301.

(5) Scott, D.E., Bayly, A.R., Abell, C., Skidmore, J. Small Molecules, Big Targets: Drug Discovery Faces the Protein–protein Interaction Challenge. Nat. Rev. Drug Discov. 2016, 15, 533–550.

(6) Pelay-Gimeno, M., Glas, A., Koch, O., Grossmann, T.N. Structure-based Design of Inhibitors of Protein–protein Interactions: Mimicking Peptide Binding Epitopes. Angew. Chem. Int. Ed. 2015, 54, 8896–8927.

(7) Chames, P., Van Regenmortel, M., Weiss, E., Baty, D. Therapeutic Antibodies: Successes, Limitations and Hopes for the Future. Br. J. Pharmacol. 2009, 157, 220–233.

(8) Ali, A.M., Atmaj, J., Van Oosterwijk, N., Groves, M.R., Dömling, A. Stapled Peptides Inhibitors: A New Window for Target Drug Discovery. Comput. Struct. Biotechnol. J. 2019, 17, 263–281.

(9) Craik, D.J., Fairlie, D.P., Liras, S., Price, D. The Future of Peptide-based Drugs. Chem. Biol. Drug. Des. 2013, 81, 136–147.

(10) Kutchukian, P.S., Yang, J.S., Verdine, G.L., Shakhnovich, E.I. All-atom Model for Stabilization of α-helical Structure in Peptides by Hydrocarbon Staples. J. Am. Chem. Soc. 2009, 131, 4622–4627.

(11) Sato, A.K., Viswanathan, M., Kent, R.B., Wood, C.R. Therapeutic Peptides: Technological Advances Driving Peptides into Development. Curr. Opin. Biotech. 2006, 17, 638–642.

(12) Klein, M. Stabilized Helical Peptides: Overview of the Technologies and its Impact on Drug Discovery. Expert. Opin. Drug. Discov. 2017, 12, 1117–1125.

(13) Walensky, L.D., Bird, G.H. Hydrocarbon-stapled Peptides: Principles, Practice, and Progress: Miniperspective. J. Med. Chem. 2014, 57, 6275–6288.

(14) Shepherd, N.E., Hoang, H.N., Abbenante, G., Fairlie, D.P. Single Turn Peptide Alpha Helices with Exceptional Stability in Water. J. Am. Chem. Soc. 2005, 127, 2974–2983.

(15) Lau, Y.H., Wu, Y., Rossmann, M., Tan, B.X., de Andrade, P., Tan, Y.S., Verma, C., McKenzie, G.J., Venkitaraman, A.R., Hyvönen, M., Spring, D.R. Double Strain-promoted Macrocyclization for the Rapid Selection of Cell-active Stapled Peptides. Angew. Chem. Int. Ed. Engl. 2015, 54, 15410–15413.

(16) Phillips, C., Roberts, L.R., Schade, M., Bazin, R., Bent, A., Davies, N.L., Moore, R., Pannifer, A.D., Pickford, A.R., Prior, S.H., Read, C.M., Scott, A., Brown, D.G., Xu, B., Irving, S.L. Design and Structure of Stapled Peptides Binding to Estrogen Receptors. J. Am. Chem. Soc. 2011, 133, 9696–9699.

(17) Kneissl, S., Loveridge, E.J., Williams, C., Crump, M.P., Allemann, R.K. Photocontrollable Peptide-based Switches Target the Anti-apoptotic Protein Bcl-xL. Chem-bioChem 2008, 9, 3046–3054.

(18) Bernal, F., Tyler, A.F., Korsmeyer, S.J., Walensky, L.D., Verdine, G.L. Reactivation of the p53 Tumor Suppressor Pathway by a Stapled p53 Peptide. J. Am. Chem. Soc. 2007, 129, 2456–2457.

(19) Walensky, L.D., Kung, A.L., Escher, I., Malia, T.J., Barbuto, S., Wright, R.D., Wagner, G., Verdine, G.L., Korsmeyer, S.J. Activation of Apoptosis in vivo by a Hydrocarbon-stapled BH3 Helix. Science 2004, 305, 1466–1470.

(20) Speltz, T.E., Fanning, S.W., Mayne, C.G., Fowler, C., Tajkhorshid, E., Greene, G.L., Moore, T.W. Stapled Peptides with γ-Methylated Hydrocarbon Chains for the Estrogen Receptor/Coactivator Interaction. Angew. Chem. Int. Ed. Engl. 2016, 55, 4252– 4255.

(21) Levine, A.J. p53, the Cellular Gatekeeper for Growth and Division. Cell 1997, 88, 323–331.

(22) Vousden, K.H., Lu, X. Live or Let Die: The Cell’s Response to p53. Nat. Rev. Cancer 2002, 2, 594–604.

(23) Momand, J., Zambetti, G.P., Olson, D.C., George, D., Levine, A.J. The mdm2 Oncogene Product Forms a Complex with the p53 Protein and Inhibits p53-Mediated Transactivation. Cell 1992, 69, 1237–1245.

(24) Eymin, B., Gazzeri, S., Brambilla, C., Brambilla, E. Mdm2 Overexpression and P14 ARF Inactivation are Two Mutually Exclusive Events in Primary Human Lung Tumors. Oncogene 2002, 21, 2750–2761.

(25) Polsky, D., Bastian, B.C., Hazan, C., Melzer, K., Pack, J., Houghton, A., Busam, K., Cordon-Cardo, C., Osman, I. HDM2 Protein Overexpression, but not Gene Amplification, is Related to Tumorigenesis of Cutaneous Melanoma. Cancer Res. 2001, 61, 7642–7646.

(26) Vassilev, L.T., Vu, B.T., Graves, B., Carvajal, D., Podlaski, F., Filipovic, Z., Kong, N., Kammlott, U., Lukacs, C., Klein, C., Fotouhi, N., Liu, E.A. In Vivo Activation of the p53 Pathway by Small-Molecule Antagonists of MDM2. Science 2004, 303, 844–848.

(27) Shangary, S., Qin, D., McEachern, D., Liu, M., Miller, R.S., Qiu, S., NikolovskaColeska, Z., Ding, K., Wang, G., Chen, J., Bernard, D., Zhang, J., Lu, Y., Gu, Q., Shah, R.B., Pienta, K.J., Ling, X., Kang, S., Guo, M., Sun, Y., Yang, D., Wang, S. Temporal Activation of p53 by a Specific MDM2 Inhibitor is Selectively Toxic to Tumors and Leads to Complete Tumor Growth Inhibition. Proc. Natl. Acad. Sci. U.S.A. 2008, 105, 3933–3938.

(28) Böttger, V., Böttger, A., Howard, S.F., Picksley, S.M., Chéne, P., GarciaEcheverria, C., Hochkeppel, H.-K., Lane, D.P. Identification of Novel mdm2 Binding Peptides by Phage Display. Oncogene 1996, 13, 2141–2147.

(29) Baek, S., Kutchukian, P.S., Verdine, G.L., Huber, R., Holak, T.A., Lee, K.W., Popowicz, G.M. Structure of the Stapled p53 Peptide Bound to Mdm2. J. Am. Chem. Soc. 2012, 134, 103–106.

(30) Chang, Y.S., Graves, B., Guerlavais, V., Tovar, C., Packman, K., To, K.-H., Olson, K.A., Kesavan, K., Gangurde, P., Mukherjee, A., Baker, T., Darlak, K., Elkin, C., Filipovic, Z., Qureshi, F.Z., Cai, H., Berry, P., Feyfant, E., Shi, X.E., Horstick, J., Annis, D.A., Manning, A.M., Fotouhi, N., Nash, H., Vassilev, L.T., Sawyer, T.K. Stapled α helical Peptide Drug Development: A Potent Dual Inhibitor of MDM2 and MDMX for p53-dependent Cancer Therapy. Proc. Natl. Acad. Sci. U.S.A. 2013, 110, E3445–E3454.

(31) Brown, C.J., Quah, S.T., Jong, J., Goh, A.M., Chiam, P.C., Khoo, K.H., Choong, M.L., Lee, M.A., Yurlova, L., Zolghadr, K., Joseph, T.L., Verma, C.S., Lane, D.P. Stapled Peptides with Improved Potency and Specificity that Activate p53. ACS Chem. Biol. 2013, 8, 506–512.

(32) Joseph, T.L., Lane, D., Verma, C.S. Stapled Peptides in the p53 Pathway: Computer Simulations Reveal Novel Interactions of the Staples With the Target Protein. Cell Cycle 2010, 9, 4560–4568.

(33) Tan, Y.S., Lane, D.P., Verma, C.S. Stapled Peptide Design: Principles and Roles of Computation. Drug Discov. Today 2016, 21, 1642–1653.

(34) Cornillie, S.P., Bruno, B.J., Lim, C.S., Cheatham III, T.E. Computational Modeling of Stapled Peptides toward a Treatment Strategy for CML and Broader Implications in the Design of Lengthy Peptide Therapeutics. J. Phys. Chem. B 2018, 122, 3864–3875.

(35) Villavicencio, B., Ligabue-Braun, R., Verli, H. All-Hydrocarbon Staples and Their Effect over Peptide Conformation under Different Force Fields. J. Chem. Inf. Model. 2018, 58, 2015–2023.

(36) Morrone, J.A., Perez, A., Deng, Q., Ha, S.N., Holloway, M.K., Sawyer, T.K., Sherborne, B.S., Brown, F.K., Dill, K.A. Molecular Simulations Identify Binding Poses and Approximate Affinities of Stapled α-helical Peptides to MDM2 and MDMX. J. Chem. Theory Comput 2017, 13, 863–869.

(37) Kussie, P.H., Gorina, S., Marechal, V., Elenbaas, B., Moreau, J., Levine, A.J., Pavletich, N.P. Structure of the MDM2 Oncoprotein Bound to the p53 Tumor Suppressor Transactivation Domain. Science 1996, 274, 948–953.

(38) Kannan, S., Partridge, A.W., Lane, D.P., Verma, C.S. The Dual Interactions of p53 with MDM2 and p300: Implications for the Design of MDM2 Inhibitors. Int. J. Mol. Sci. 2019, 20, 5996.

(39) Yadahalli, S., Neira, J.L., Johnson, C.M., Tan, Y.S., Rowling, P.J., Chattopadhyay, A., Verma, C.S., Itzhaki, L.S. Kinetic and Thermodynamic Effects of Phosphorylation on p53 Binding to MDM2. Sci. Rep. 2019, 9, 1–15.

(40) Mukherjee, A., Bagchi, B. Correlation between Rate of Folding, Energy Landscape, and Topology in the Folding of a Model Protein HP-36. J. Chem. Phys. 2003, 118, 4733–4747.

(41) Roy, S., Bagchi, B. Comparative Study of Protein Unfolding in Aqueous Urea and Dimethyl Sulfoxide Solutions: Surface Polarity, Solvent Specificity, and Sequence of Secondary Structure Melting. J. Phys. Chem. B 2014, 118, 5691–5697.

(42) Caballero-Herrera, A., Nordstrand, K., Berndt, K.D., Nilsson, L. Effect of Urea on Peptide Conformation in Water: Molecular Dynamics and Experimental Characterization. Biophys. J. 2005, 89, 842–857.

(43) Sarma, R., Paul, S. Interactions of S-peptide Analogue in Aqueous Urea and Trimethylamine-N-oxide Solutions: A Molecular Dynamics Simulation Study. J. Chem. Phys. 2013, 139, 07B614 1.

(44) Ang, H.C., Joerger, A.C., Mayer, S., Fersht, A.R. Effects of Common Cancer Mutations on Stability and DNA Binding of Full-length p53 Compared with Isolated Core Domains. J. Biol. Chem. 2006, 281, 21934–21941.

(45) Bullock, A.N., Fersht, A.R. Rescuing the Function of Mutant p53. Nat. Rev. Cancer 2001, 1, 68–76.

(46) Best, R.B., Zhu, X., Shim, J., Lopes, P.E., Mittal, J., Feig, M., MacKerell Jr, A.D. Optimization of the Additive CHARMM All-Atom Protein Force Field Targeting Improved Sampling of The Backbone φ, ψ and Side-Chain χ1 and χ2 Dihedral Angles. J. Chem. Theory Comput. 2012, 8, 3257–3273.

(47) Vanommeslaeghe, K., Hatcher, E., Acharya, C., Kundu, S., Zhong, S., Shim, J., Darian, E., Guvench, O., Lopes, P., Vorobyov, I., Mackerell Jr., A.D. CHARMM General Force Field: A Force Field for Drug-Like Molecules Compatible with the CHARMM All-Atom Additive Biological Force Fields. J. Comput. Chem. 2010, 31, 671–690.

(48) Dastidar, S.G., Lane, D.P., Verma, C.S. Modulation of p53 Binding to MDM2: Computational Studies Reveal Important Roles of Tyr100. BMC Bioinformatics. 2009; pp 1–11.

(49) Speltz, T.E., Mayne, C.G., Fanning, S.W., Siddiqui, Z., Tajkhorshid, E., Greene, G.L., Moore, T.W. A “Cross-Stitched” Peptide with Improved Helicity and Proteolytic Stability. Org. Biomol. Chem. 2018, 16, 3702–3706.

(50) Mark, P., Nilsson, L. Structure and Dynamics of the TIP3P, SPC, and SPC/E Water Models at 298 K. J. Phys. Chem. A 2001, 105, 9954–9960.

(51) Petrova, S.S., Solov’ev, A.D. The Origin of the Method of Steepest Descent. Hist. Math. 1997, 24, 361–375.

(52) Bussi, G., Donadio, D., Parrinello, M. Canonical Sampling through Velocity Rescaling. J. Chem. Phys. 2007, 126, 014101.

(53) Parrinello, M., Rahman, A. Polymorphic Transitions in Single Crystals: A New Molecular Dynamics Method. J. Appl. Phys. 1981, 52, 7182–7190.

(54) Van Der Spoel, D., Lindahl, E., Hess, B., Groenhof, G., Mark, A.E., Berendsen, H.J. GROMACS: Fast, Flexible, and Free. J. Comput. Chem. 2005, 26, 1701–1718.

(55) Allen, M.P., Tildesley, D.J. Computer Simulation of Liquids; Oxford university press, 2017.

(56) Andersen, H.C. Rattle: A “Velocity” Version of the Shake Algorithm for Molecular Dynamics Calculations. J. Comput. Phys. 1983, 52, 24–34.

(57) Darden, T., York, D., Pedersen, L. Particle Mesh Ewald: An Nlog (N) Method for Ewald Sums in Large Systems. J. Chem. Phys. 1993, 98, 10089–10092.

(58) Humphrey, W., Dalke, A., Schulten, K.VMD: Visual Molecular Dynamics. J. Mol. Graph. 1996, 14, 33–38.

(59) Genheden, S., Ryde, U. The MM/PBSA and MM/GBSA Methods to Estimate Ligandbinding Affinities. Expert Opin. Drug Discov. 2015, 10, 449–461.

(60) Gilson, M.K., Given, J.A., Bush, B.L., McCammon, J.A. The Statistical-thermodynamic Basis for Computation of Binding Affinities: A Critical Review. Biophys. J. 1997, 72, 1047–1069.

(61) Stoica, I., Sadiq, S.K., Coveney, P.V. Rapid and Accurate Prediction of Binding Free Energies for Saquinavir-bound HIV-1 Proteases. J. Am. Chem. Soc. 2008, 130, 2639–2648.

(62) Sadiq, S.K., Wright, D.W., Kenway, O.A., Coveney, P.V. Accurate Ensemble Molecular Dynamics Binding Free Energy Ranking of Multidrug-resistant HIV-1 Proteases. J Chem Inf Model 2010, 50, 890–905.

(63) Im, W., Lee, M.S., Brooks III, C.L. Generalized Born Model with a Simple Smoothing Function. J. Comput. Chem. 2003, 24, 1691–1702.

(64) Ahalawat, N., Mondal, J. Assessment and Optimization of Collective Variables for Protein Conformational Landscape: GB1 β-Hairpin as a Case Study. J. Chem. Phys 2018, 149, 094101.

(65) Xu, J., Huang, L., Shakhnovich, E.I. The Ensemble Folding Kinetics of the FBP28 WW Domain Revealed by an All-atom Monte Carlo Simulation in a Knowledge-based Potential. Proteins 2011, 79, 1704–1714.

(66) Ma, W., Whitley, K.D., Chemla, Y.R., Luthey-Schulten, Z., Schulten, K. Free-energy Simulations Reveal Molecular Mechanism for Functional Switch of a DNA Helicase. Elife 2018, 7, e34186.

(67) Sarkar, S., Maity, A., Sarma Phukon, A., Ghosh, S., Chakrabarti, R. Salt Induced Structural Collapse, Swelling, and Signature of Aggregation of Two SsDNA Strands: Insights from Molecular Dynamics Simulation. J. Phys. Chem. B 2019, 123, 47–56.

(68) Maity, A., Sarkar, S., Theeyancheri, L., Chakrabarti, R. Choline Chloride as a Nano-Crowder Protects HP-36 from Urea-Induced Denaturation: Insights from Solvent Dynamics and Protein-Solvent Interactions. ChemPhysChem 2020, 21, 552–567.

(69) Gao, Y., Li, Y., Mou, L., Lin, B., Zhang, J.Z., Mei, Y. Correct Folding of an α-helix and a β-hairpin Using a Polarized 2D Torsional Potential. Sci. Rep. 2015, 5, 10359.

(70) Bonomi, M., Branduardi, D., Bussi, G., Camilloni, C., Provasi, D., Raiteri, P., Donadio, D., Marinelli, F., Pietrucci, F., Broglia, R.A., Parrinello, M. PLUMED: A Portable Plugin for Free-energy Calculations with Molecular Dynamics. Comput. Phys. Commun. 2009, 180, 1961–1972.

(71) Pietrucci, F., Laio, A. A Collective Variable for the Efficient Exploration of Protein Beta-sheet Structures: Application to SH3 and GB1. J. Chem. Theory Comput. 2009, 5, 2197–2201.

(72) Zondlo, S.C., Lee, A.E., Zondlo, N.J. Determinants of Specificity of MDM2 for the Activation Domains of p53 and p65: Proline27 Disrupts the MDM2-binding Motif of p53. Biochemistry 2006, 45, 11945–11957.

(73) Xiong, K., Zwier, M.C., Myshakina, N.S., Burger, V.M., Asher, S.A., Chong, L.T. Direct Observations of Conformational Distributions of Intrinsically Disordered p53 Peptides Using UV Raman and Explicit Solvent Simulations. J. Phys. Chem. A 2011, 115, 9520–9527.

(74) Guo, Z., Mohanty, U., Noehre, J., Sawyer, T.K., Sherman, W., Krilov, G. Probing the α-helical Structural Stability of Stapled p53 Peptides: Molecular Dynamics Simulations and Analysis. Chem. Biol. Drug Des. 2010, 75, 348–359.

(75) Sim, A.Y., Joseph, T., Lane, D.P., Verma, C. Mechanism of Stapled Peptide Binding to MDM2: Possible Consequences for Peptide Design. J. Chem. Theory Comput. 2014, 10, 1753–1761.

(76) Valiente, P.A., Becerra, D., Kim, P.M. A Method to Calculate the Relative Binding Free Energy Differences of α-Helical Stapled Peptides. J. Org. Chem. 2020, 85, 1644– 1651.

(77) Greenidge, P.A., Kramer, C., Mozziconacci, J.-C., Wolf, R.M. MM/GBSA Binding Energy Prediction on the PDBbind Data Set: Successes, Failures, and Directions for Further Improvement. J. Chem. Inf. Model. 2013, 53, 201–209.

(78) Chen, J., Wang, J., Xu, B., Zhu, W., Li, G. Insight into Mechanism of Small Molecule Inhibitors of the MDM2–p53 Interaction: Molecular Dynamics Simulation and Free Energy Analysis. J. Mol. Graph. Model. 2011, 30, 46–53.

(79) Chen, J., Wang, J., Zhang, Q., Chen, K., Zhu, W. Probing Origin of Binding Difference of Inhibitors to MDM2 and MDMX by Polarizable Molecular Dynamics Simulation and QM/MM-GBSA Calculation. Sci. Rep. 2015, 5, 17421.

(80) Pazgier, M., Liu, M., Zou, G., Yuan, W., Li, C., Li, C., Li, J., Monbo, J., Zella, D., Tarasov, S.G., et al. Structural Basis for High-affinity Peptide Inhibition of p53 Interactions with MDM2 and MDMX. Proc. Natl. Acad. Sci. U.S.A. 2009, 106, 4665–4670.

(81) Chi, S.-W., Lee, S.-H., Kim, D.-H., Ahn, M.-J., Kim, J.-S., Woo, J.-Y., Torizawa, T., Kainosho, M., Han, K.-H. Structural Details on mdm2-p53 Interaction. J. Biol. Chem. 2005, 280, 38795–38802.

(82) Böttger, A., Böttger, V., Garcia-Echeverria, C., Chéne, P., Hochkeppel, H.-K., Sampson, W., Ang, K., Howard, S.F., Picksley, S.M., Lane, D.P. Molecular Characterization of the hdm2-p53 Interaction. J. Mol. Biol. 1997, 269, 744–756.

(83) Schon, O., Friedler, A., Bycroft, M., Freund, S.M., Fersht, A.R. Molecular Mechanism of the Interaction between MDM2 and p53. J. Mol. Biol. 2002, 323, 491–501.

(84) Zhou, G., Pantelopulos, G.A., Mukherjee, S., Voelz, V.A. Bridging Microscopic and Macroscopic Mechanisms of p53-MDM2 Binding with Kinetic Network Models. Biophys. J. 2017, 113, 785–793.

(85) Paul, F., Wehmeyer, C., Abualrous, E.T., Wu, H., Crabtree, M.D., Schöneberg, J., Clarke, J., Freund, C., Weikl, T.R., Noé, F. Protein-peptide Association Kinetics Beyond the Seconds Timescale from Atomistic Simulations. Nat. Commun. 2017, 8, 1–10.

(86) Best, R.B., Zheng, W., Mittal, J. Balanced Protein–water Interactions Improve Properties of Disordered Proteins and Non-specific Protein Association. J. Chem. Theory Comput. 2014, 10, 5113–5124.

(87) Boehr, D.D., Nussinov, R., Wright, P.E. The Role of Dynamic Conformational Ensembles in Biomolecular Recognition. Nat. Chem. Biol. 2009, 5, 789–796.

(88) Chodera, J.D., Mobley, D.L. Entropy-enthalpy Compensation: Role and Ramifications in Biomolecular Ligand Recognition and Design. Annu. Rev. Biophys. 2013, 42, 121– 142.

(89) Gimeno, A., Delgado, S., Valverde, P., Bertuzzi, S., Berbís, M.A., Echavarren, J., Lacetera, A., Martín-Santamaría, S., Surolia, A., Ca ñada, F.J., et al. Minimizing the Entropy Penalty for Ligand Binding: Lessons from the Molecular Recognition of the Histo Blood-group Antigens by Human Galectin-3. Angew Chem Int Ed Engl 2019, 58, 7268–7272.

(90) Chang, C.E. A., Chen, W., Gilson, M.K. Ligand Configurational Entropy and Protein Binding. Proc. Natl. Acad. Sci. U.S.A. 2007, 104, 1534–1539.

(91) Li, Y., Cong, Y., Feng, G., Zhong, S., Zhang, J.Z., Sun, H., Duan, L. The Impact of Interior Dielectric Constant and Entropic Change on HIV-1 Complex Binding Free Energy Prediction. Struct. Dyn. 2018, 5, 064101.

(92) Hou, T., Wang, J., Li, Y., Wang, W. Assessing the Performance of the MM/PBSA and MM/GBSA Methods. 1. The Accuracy of Binding Free Energy Calculations Based on Molecular Dynamics Simulations. J. Chem. Inf. Model 2011, 51, 69–82.

