## Supporting Information for "Effect of Stapling on the Thermodynamics of mdm2-p53 Binding"

Atanu Maity[†], Asha Rani Choudhury[†] and Rajarshi Chakrabarti[*]
Department of Chemistry, Indian Institute of Technology Bombay, Mumbai, Powai 400076

*

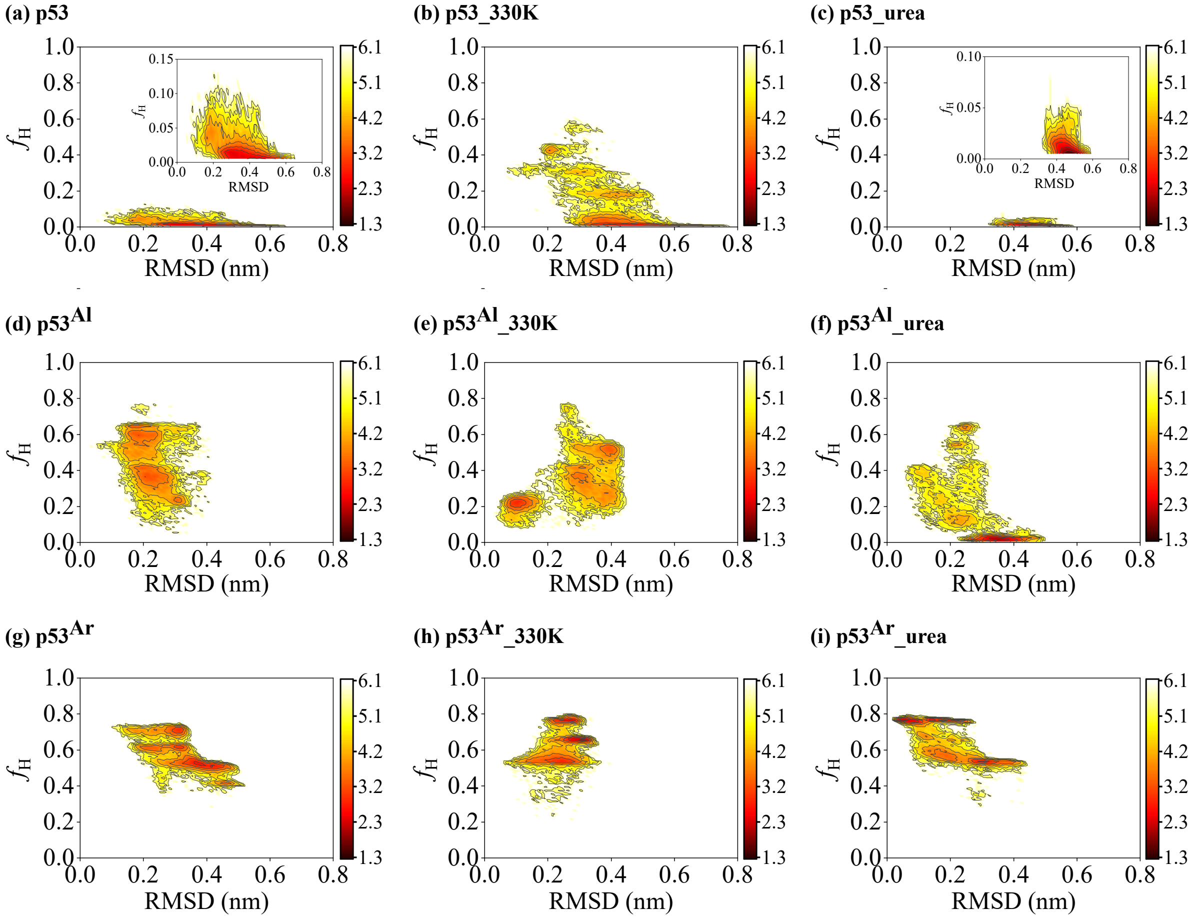


Figure S1: 2D histogram of RMSD (Root mean square deviation)-Vs-helical fraction plot for the p53 peptide and the p53 peptides with aliphatic and aromatic staples in different conditions (i.e. in the aqueous medium, at high temperature, and in the presence of urea) showing variation in the conformation in the presence of stapling agents. The dark red-to-yellow-to-whitee color scale represents increasing conformational free energy in kcal/mol.

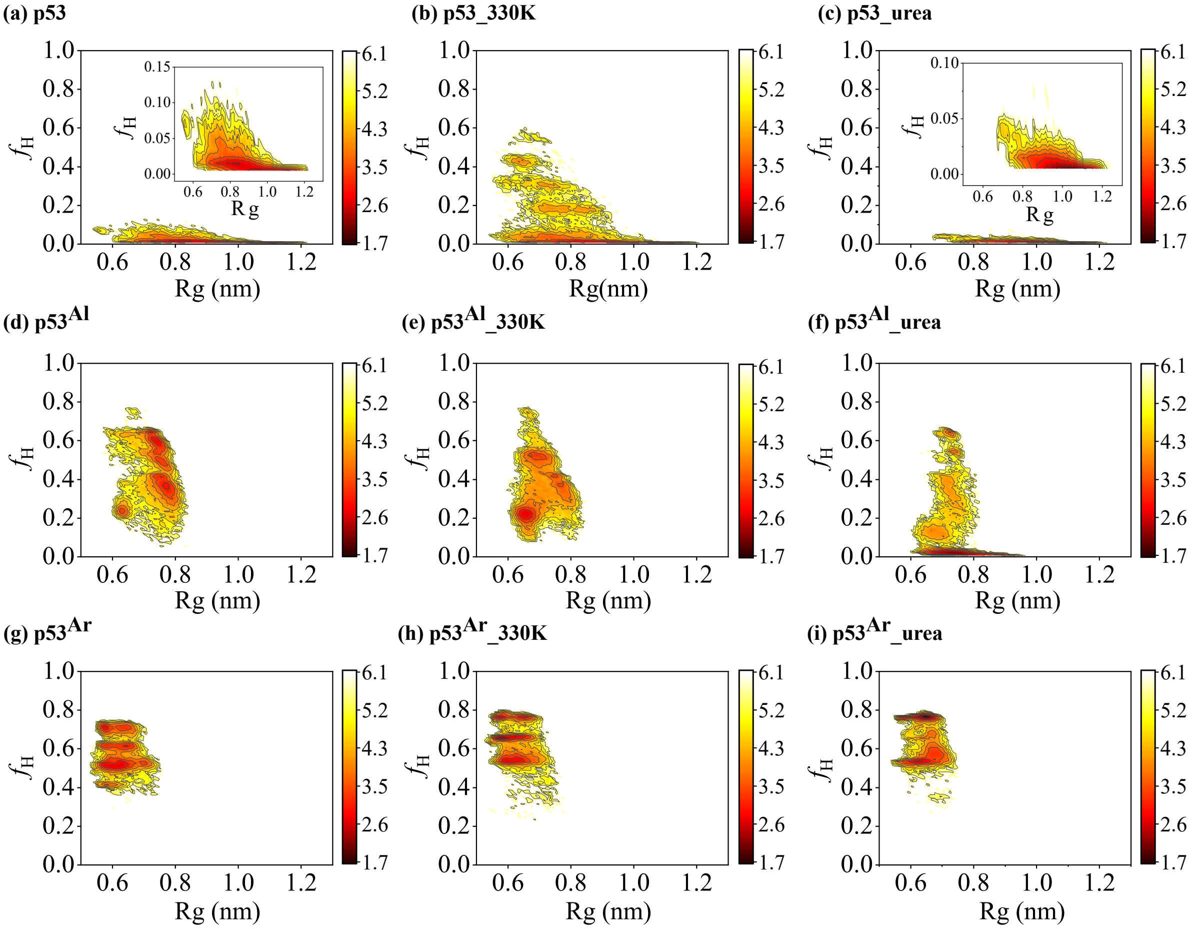


Figure S2: 2D histogram of $R_g$ (Radius of gyration)-Vs-helical fraction plot for the p53 peptide and the p53 peptides with aliphatic and aromatic staples in different conditions (i.e. in the aqueous medium, at high temperature, and in the presence of urea) showing variation in the conformation in the presence of stapling agents. The dark red-to-yellow-to-white color scale represents increasing conformational free energy in kcal/mol.

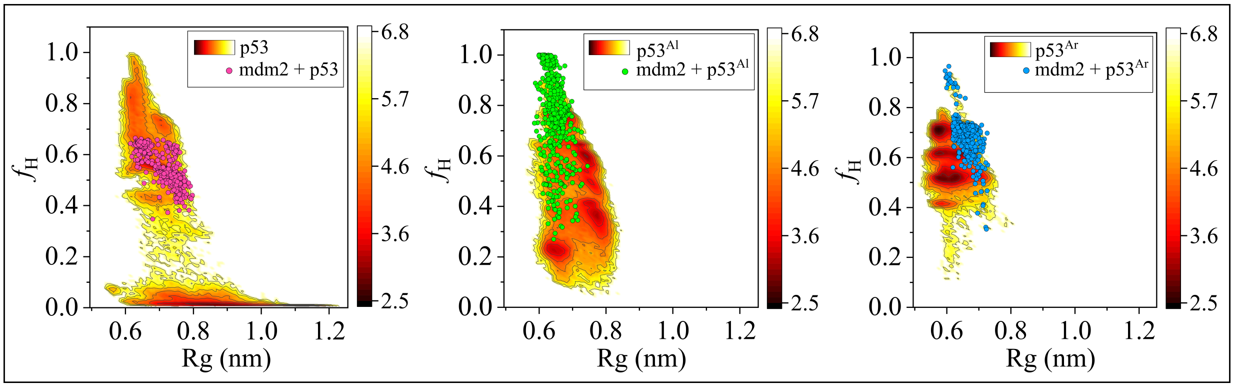


Figure S3: Quantification of unfolded to folded conformational transition during complexation. The $R_g$ and helicity of the conformations of (a) p53+mdm2, (b) mdm2 + p53$^{Al}$, and (c) mdm2 + p53$^{Ar}$ system are plotted on the $R_g$-Vs-helicity free energy surface of p53, p53$^{Al}$ and p53$^{Ar}$ respectively. There is large energy gap between the energy minimum of p53 and points corresponding to p53+mdm2 (magenta circles) whereas there is a considerable overlap between the energy minima for p53$^{Al}$ and p53$^{Ar}$ with the points corresponding to p53$^{Al}$ (green circles) and p53$^{Ar}$ (cyan circles) respectively.

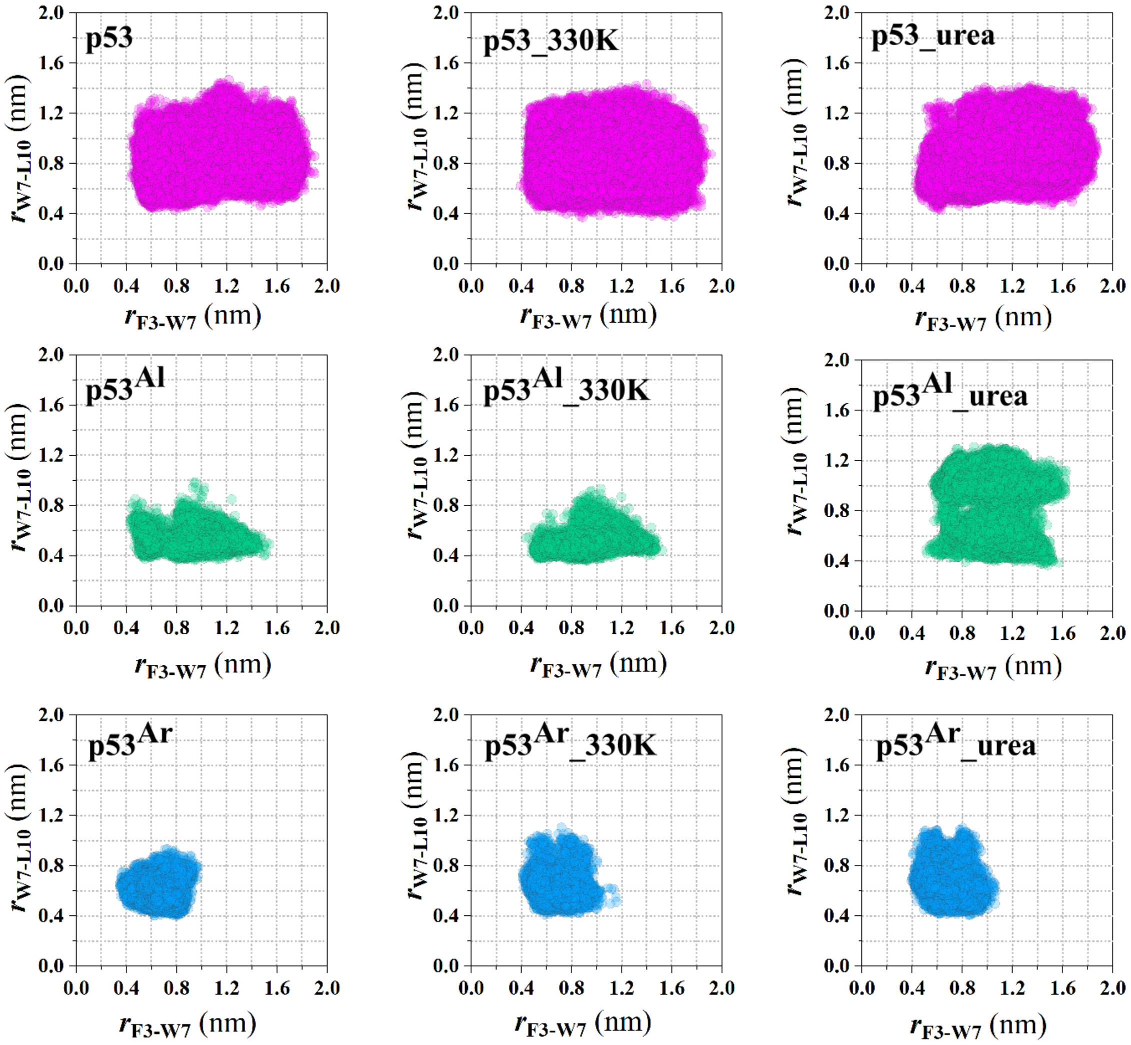


Figure S4: Plot of $r_{F3-W7}$-vs-$r_{W7-L10}$ for p53, p53$^{\mathrm{Al}}$, and p53$^{\mathrm{Ar}}$ in different conditions

Table S1: Energy components for the three simulations of **mdm2**. All energy values and standard errors are in kcal/mol.

| Energy components | mdm2 (simulation 1) | mdm2 (simulation 2) | mdm2 (simulation 3) | Average |
|---|---|---|---|---|
| $E_{Elec}$ | -1550.90 ±203.51 | -1607.59 ±198.50 | -1574.90 ±198.22 | -1577.80 ±23.23 |
| $E_{Vdw}$ | -296.42 ±15.91 | -285.81 ±15.43 | -287.13 ±15.19 | -289.79 ±4.72 |
| $E_{Internal}$ | 1986.60 ±29.15 | 1981.54 ±29.12 | 1982.92 ±28.94 | 1983.69 ±2.14 |
| $E_{Solv(polar)}$ | -1558.44 ±198.94 | -1501.07 ±194.03 | -1535.24 ±193.27 | -1531.58 ±23.56 |
| $E_{Solv(non-polar)}$ | 33.06 ±0.67 | 33.37 ±0.63 | 33.58 ±0.59 | 33.34 ±0.21 |
| $E_{Elec+Solv(polar)}$ | -3109.34 ±16.47 | -3108.66 ±15.94 | -3110.98 ±16.25 | -3109.38 ±0.6 |
| $H$ | -1386.10 ±28.74 | -1379.56 ±28.45 | -1380.77 ±27.85 | -1382.14 ±2.84 |
| $-TS$ | -355.40 ±0.53 | -369.71 ±0.17 | -378.14 ±0.36 | -367.75 ±9.39 |

Table S2: Energy components for the three simulations of **mdm2+p53**. All energy values and standard errors are in kcal/mol.

| Energy components | mdm2+p53 (simulation 1) | mdm2+p53 (simulation 2) | mdm2+p53 (simulation 3) | Average |
|---|---|---|---|---|
| $E_{Elec}$ | -1868.45 ±223.07 | -1765.45 ±227.80 | -1971.43 ±221.30 | -1868.44 ±84.09 |
| $E_{Vdw}$ | -364.14 ±16.75 | -358.39 ±17.02 | -357.21 ±17.42 | -359.91 ±3.03 |
| $E_{Internal}$ | 2303.48 ±31.10 | 2305.99 ±31.36 | 2310.67 ±30.93 | 2306.71 ±2.98 |
| $E_{Solv(polar)}$ | -1732.36 ±218.20 | -1842.15 ±222.23 | -1644.16 ±216.22 | -1739.56 ±80.99 |
| $E_{Solv(non-polar)}$ | 37.17 ±0.82 | 37.76 ±0.83 | 37.48 ±0.81 | 37.47 ±0.24 |
| $E_{Elec+Solv(polar)}$ | -3600.81 ±17.77 | -3607.60 ±18.0 | -3615.59 ±17.48 | -3608.00 ±6.04 |
| $H$ | -1624.29 ±30.66 | -1622.24 ±31.37 | -1624.65 ±30.38 | -1623.73 ±1.06 |
| $-TS$ | -429.40 ±1.13 | -431.40 ±2.69 | -427.55 ±0.91 | -429.45 ±1.57 |

Table S3: Energy components for the three simulations of $\mathbf{mdm2 + p53^{Al}}$. All energy values and standard errors are in kcal/mol.

| Energy components | mdm2 + p53$^{Al}$ (simulation 1) | mdm2 + p53$^{Al}$ (simulation 2) | mdm2 + p53$^{Al}$ (simulation 3) | Average |
|---|---|---|---|---|
| $E_{Elec}$ | -1866.26 ±227.91 | -1943.48 ±229.18 | -1920.53 ±218.78 | -1910.09 ±32.38 |
| $E_{Vdw}$ | -374.87 ±17.53 | -374.46 ±17.02 | -372.92 ±17.23 | -374.08 ±0.84 |
| $E_{Internal}$ | 2345.72 ±31.39 | 2341.94 ±31.67 | 2341.03 ±31.51 | 2342.90 ±2.03 |
| $E_{Solv(polar)}$ | -1765.86 ±223.54 | -1683.43 ±223.50 | -1709.17 ±213.80 | -1719.49 ±34.43 |
| $E_{Solv(non-polar)}$ | 37.18 ±0.76 | 37.11 ±0.89 | 37.20 ±0.81 | 37.17 ±0.04 |
| $E_{Elec+Solv(polar)}$ | -3632.12 ±17.57 | -3626.91 ±18.41 | -3629.70 ±17.70 | -3629.58 ±2.13 |
| $H$ | -1624.09 ±31.73 | -1622.31 ±31.85 | -1624.39 ±30.96 | -1623.60 ±0.92 |
| $-TS$ | -423.68 ±0.18 | -432.45 ±0.48 | -425.12 ±0.17 | -427.08 ±3.84 |

Table S4: Energy components for the three simulations of $\mathbf{mdm2 + p53^{Ar}}$. All energy values and standard errors are in kcal/mol.

| Energy components | mdm2 + p53$^{Ar}$ (simulation 1) | mdm2 + p53$^{Ar}$ (simulation 2) | mdm2 + p53$^{Ar}$ (simulation 3) | Average |
|---|---|---|---|---|
| $E_{Elec}$ | -1903.94 ±224.35 | -1972.60 ±218.79 | -1955.18 ±222.07 | -1943.91 ±29.14 |
| $E_{Vdw}$ | -367.50 ±17.75 | -362.22 ±16.22 | -364.20 ±17.43 | -364.64 ±2.18 |
| $E_{Internal}$ | 2324.47 ±31.66 | 2324.41 ±31.23 | 2327.33 ±31.35 | 2325.40 ±1.36 |
| $E_{Solv(polar)}$ | -1725.82 ±219.08 | -1661.12 ±214.53 | -1677.61 ±217.47 | -1688.18 ±27.45 |
| $E_{Solv(non-polar)}$ | 36.97 ±0.88 | 37.16 ±0.73 | 37.01 ±0.82 | 37.05 ±0.08 |
| $E_{Elec+Solv(polar)}$ | -3629.76 ±18.29 | -3633.72 ±18.39 | -3632.79 ±17.81 | -3632.09 ±1.69 |
| $H$ | -1635.83 ±31.28 | -1634.36 ±31.59 | -1632.65 ±31.00 | -1634.28 ±1.30 |
| $-TS$ | -429.89 ±1.12 | -431.72 ±1.40 | -417.48 ±0.48 | -426.36 ±6.32 |

Table S5: Energy components for the three simulations of **p53**. All energy values and standard errors are in kcal/mol.

| Energy components | p53 (simulation 1) | p53 (simulation 2) | p53 (simulation 3) | Average |
|---|---|---|---|---|
| $E_{Elec}$ | -110.78 ±51.21 | -113.94 ±47.87 | -118.82 ±53.20 | -114.51 ±3.31 |
| $E_{Vdw}$ | 3.06 ±6.16 | 1.17 ±8.60 | -1.34 ±8.52 | 0.96 ±1.80 |
| $E_{Internal}$ | 309.50 ±13.48 | 310.67 ±11.88 | 311.52 ±11.74 | 310.57 ±0.83 |
| $E_{Solv(polar)}$ | -387.99 ±48.66 | -385.21 ±45.41 | -381.25 ±49.39 | -384.81 ±2.76 |
| $E_{Solv(non-polar)}$ | 8.69 ±0.35 | 8.54 ±0.53 | 8.42 ±0.56 | 8.55 ±0.11 |
| $E_{Elec+Solv(polar)}$ | -498.76 ±6.56 | -499.15 ±6.53 | -500.07 ±7.76 | -499.32 ±0.55 |
| $H$ | -177.51 ±14.20 | -178.76 ±13.34 | -181.47 ±14.59 | -179.25 ±1.65 |
| $-TS$ | -114.02 ±0.07 | -116.63 ±0.54 | -116.09 ±0.11 | -115.58 ±1.12 |

Table S6: Energy components for the three simulations of $\mathbf{p53^{Al}}$. All energy values and standard errors are in kcal/mol.

| Energy components | $p53^{Al}$ (simulation 1) | $p53^{Al}$ (simulation 2) | $p53^{Al}$ (simulation 3) | Average |
|---|---|---|---|---|
| $E_{Elec}$ | -188.58 ±49.43 | -165.13 ±40.75 | -227.12 ±41.20 | -193.61 ±25.56 |
| $E_{Vdw}$ | -15.63 ±5.68 | -14.08 ±5.66 | -15.74 ±5.60 | -15.15 ±0.76 |
| $E_{Internal}$ | 344.45 ±15.19 | 345.87 ±12.41 | 348.08 ±12.23 | 346.14 ±1.49 |
| $E_{Solv(polar)}$ | -335.37 ±45.76 | -354.60 ±39.30 | -301.30 ±38.43 | -330.42 ±22.04 |
| $E_{Solv(non-polar)}$ | 8.10 ±0.26 | 8.32 ±0.24 | 7.93 ±0.22 | 8.12 ±0.16 |
| $E_{Elec+Solv(polar)}$ | -523.95 ±7.53 | -519.73 ±6.97 | -528.41 ±6.69 | -524.03 ±3.54 |
| $H$ | -187.03 ±15.35 | -179.62 ±12.34 | -188.14 ±11.91 | -184.93 ±3.78 |
| $-TS$ | -86.38 ±0.43 | -91.53 ±0.84 | -79.31 ±0.17 | -85.74 ±5.01 |

Table S7: Energy components for the three simulations of $\mathbf{p53^{Ar}}$. All energy values and standard errors are in kcal/mol.

| Energy components | $p53^{Ar}$ (simulation 1) | $p53^{Ar}$ (simulation 2) | $p53^{Ar}$ (simulation 3) | Average |
|---|---|---|---|---|
| $E_{Elec}$ | -183.83 ±38.21 | -183.10 ±38.54 | -188.74 ±38.37 | -185.22 ±2.50 |
| $E_{Vdw}$ | -7.04 ±7.14 | -7.65 ±7.22 | -8.65 ±6.80 | -7.78 ±0.66 |
| $E_{Internal}$ | 330.74 ±12.01 | 333.37 ±11.93 | 333.52 ±11.88 | 332.68 ±1.38 |
| $E_{Solv(polar)}$ | -345.11 ±36.04 | -346.99 ±36.59 | -342.13 ±36.80 | -344.80 ±1.92 |
| $E_{Solv(non-polar)}$ | 7.67 ±0.32 | 7.69 ±0.30 | 7.56 ±0.30 | 7.64 ±0.06 |
| $E_{Elec+Solv(polar)}$ | -528.94 ±6.66 | -530.09 ±6.37 | -531.05 ±6.05 | -530.03 ±0.86 |
| $H$ | -197.58 ±11.91 | -196.27 ±12.36 | -198.61 ±12.01 | -197.49 ±0.96 |
| $-TS$ | -77.29 ±0.63 | -72.60 ±0.28 | -73.82 ±0.21 | -74.57 ±1.99 |

Table S8: Binding free energy of **mdm2-p53 binding** from simulation 1. All energy values are in kcal/mol.

| Energy components | mdm2+p53 | mdm2 | p53 | $\Delta E$ |
|---|---|---|---|---|
| $E_{Elec}$ | -1868.44 ±84.09 | -1577.80 ±23.23 | -114.51 ±3.31 | -176.14 |
| $E_{Vdw}$ | -359.91 ±3.03 | -289.79 ±4.72 | 0.96 ±1.80 | -71.09 |
| $E_{Internal}$ | 2306.71 ±2.98 | 1983.69 ±2.14 | 310.57 ±0.83 | 12.46 |
| $E_{Solv(polar)}$ | -1739.56 ±80.99 | -1531.58 ±23.56 | -384.81 ±2.76 | 176.84 |
| $E_{Solv(non-polar)}$ | 37.47 ±0.24 | 33.34 ±0.21 | 8.55 ±0.11 | -4.41 |
| $E_{Elec+Solv(polar)}$ | -3608.00 ±6.04 | -3109.38 ±0.60 | -499.32 ±0.55 | 0.70 |
| $H$ | -1623.73 ±1.06 | -1382.14 ±2.84 | -179.26 ±1.65 | -62.34 |
| $-TS$ | -429.45 ±1.57 | -367.75 ±9.39 | -115.58 ±1.39 | 53.88 |
| $G$ | | | | -8.46 ±0.69 |

Table S9: Binding free energy of $\mathbf{mdm2+p53^{Al}}$ **binding** from simulation 1. All energy values are in kcal/mol.

| Energy components | $mdm2+p53^{Al}$ | mdm2 | $p53^{Al}$ | $\Delta E$ |
|---|---|---|---|---|
| $E_{Elec}$ | -1910.91 ±32.38 | -1577.80 ±23.23 | -193.61 ±25.56 | -138.69 |
| $E_{Vdw}$ | -374.08 ±0.84 | -289.79 ±4.72 | -15.15 ±0.76 | -69.14 |
| $E_{Internal}$ | 2342.90 ±2.03 | 1983.69 ±2.14 | 346.14 ±1.49 | 13.07 |
| $E_{Solv(polar)}$ | -1719.49 ±34.43 | -1531.58 ±23.56 | -330.42 ±22.04 | 142.52 |
| $E_{Solv(non-polar)}$ | 37.17 ±0.04 | 33.34 ±0.21 | 8.12 ±0.16 | -4.29 |
| $E_{Elec+Solv(polar)}$ | -3629.58 ±2.13 | -3109.38 ±0.60 | -524.03 ±3.54 | 3.83 |
| $H$ | -1623.60 ±0.92 | -1382.14 ±2.84 | -184.93 ±3.78 | -56.53 |
| $-TS$ | -427.08 ±3.84 | -367.75 ±9.39 | -85.74 ±5.01 | 26.41 |
| $G$ | | | | -30.12 ±2.97 |

Table S10: Binding free energy of **mdm2+p53$^{\mathbf{Ar}}$ binding** from simulation 1. All energy values are in kcal/mol.

| Energy components | mdm2+p53$^{Ar}$ | mdm2 | p53$^{Ar}$ | $\Delta E$ |
|---|---|---|---|---|
| $E_{Elec}$ | -1943.91 ±29.14 | -1577.80 ±23.23 | -185.22 ±2.50 | -180.89 |
| $E_{Vdw}$ | -364.64 ±2.18 | -289.79 ±4.72 | -7.78 ±0.66 | -67.07 |
| $E_{Internal}$ | 2325.40 ±1.36 | 1983.69 ±2.14 | 332.68 ±1.38 | 9.04 |
| $E_{Solv(polar)}$ | -1688.18 ±27.45 | -1531.58 ±23.56 | -344.80 ±1.92 | 188.20 |
| $E_{Solv(non-polar)}$ | 37.05 ±0.08 | 33.34 ±0.21 | 7.64 ±0.06 | -3.93 |
| $E_{Elec+Solv(polar)}$ | -3632.09 ±1.69 | -3109.38 ±0.60 | -530.03 ±0.86 | 7.32 |
| $H$ | -1634.28 ±1.30 | -1382.14 ±2.84 | -197.49 ±0.96 | -54.65 |
| $-TS$ | -426.36 ±6.32 | -367.75 ±9.39 | -74.57 ±1.99 | 15.95 |
| $G$ | | | | -38.70 ±2.46 |
